# Thermodynamic analog of integrate-and-fire neuronal networks by maximum entropy modelling

**DOI:** 10.1101/2024.01.18.576167

**Authors:** T. S. A. N. Simões, C. I. N. Sampaio Filho, H. J. Herrmann, J. S. Andrade, L. de Arcangelis

## Abstract

Relying on maximum entropy arguments, certain aspects of time-averaged experimental neuronal data have been recently described using Ising-like models, allowing the study of neuronal networks under an analogous thermodynamical framework. Here, we apply for the first time the Maximum Entropy method to an Integrate-and-fire (IF) model that can be tuned at criticality, offering a controlled setting for a systematic study of criticality and finite-size effects in spontaneous neuronal activity, as opposed to experiments. We show that generalized Ising models that accurately predict the average local activities and correlation functions between neurons of the IF model networks in the critical state exhibit a spin glass phase for low temperatures, having mostly negative intrinsic fields and a bimodal distribution of interaction constants that tends to become unimodal for larger networks. Results appear to be affected by sample-to-sample connectivity variations and subsampling. Furthermore, we also found that networks with higher percentage of inhibitory neurons lead to Ising-like systems with reduced thermal fluctuations. Finally, considering only neuronal pairs associated with the largest correlation functions allows the study of larger system sizes.

**Author summary:** Brain activity, either stimulated or spontaneous, *in vivo* or *in vitro*, exhibits complex spatiotemporal behavior. Trying to make sense of it, several research groups have analyzed time-averaged experimental neuronal data using maximum entropy arguments, mapping the neuronal dynamics into a generalized Ising-like model and allowing to study neuronal data using tools typical of critical phenomena. However, the intricacy of real biological networks in experimental settings pose challenges in the precision and reliability of the neuronal measurements. Here, we apply for the first time the Maximum Entropy Method to an Integrate-and-fire model with synaptic plasticity, providing a foundation for a more systematic and comprehensive study of spontaneous brain activity. We show that generalized Ising models are able to reproduce the numerical time-averaged data of local activities and correlation functions of integrate-and-fire neurons and predict qualitatively higher-order quantities such as the three-point correlation functions across triplets of neurons. We show that subsampling affects the efficiency of the mapping and that the analogous thermodynamics functions of the Ising-like models depend on sample-to-sample network variations and on the presence of inhibition in the neural network.

## Introduction

Biological neural networks are highly complex systems [1], due to the large number of interacting degrees of freedom and the connectivity properties of its constituent elements. Neurons interact by generating action potentials (“firing” or “spiking”), with a biophysical mechanism first explained quantitatively by Hodgkin and Huxley in 1952 [2]. Their dynamics can be characterized by discretized time series of these firing patterns in binary notation [3, 4]. This description suggests an analogy with the binary spin states in Ising models. Indeed, Ising-like descriptions of brain activity can be traced back to fifty years ago, to the work by Little [5] and Hopfield [6], followed by the work of Amit et al. [7], where the authors used a spin glass model to describe neural networks. More recently [8–15], generalized Ising models have been used to describe the dynamics of recordings from stimulated neuronal activity, using the so-called Maximum Entropy Modelling (MEM) method [16]. The method consists in finding the least biased (or maximum entropy) probability distribution that is consistent with a given set of statistical measurements from the system under consideration but otherwise imposes no further constraints [17]. When this method is applied to describe the individual firing rates and correlation functions between neurons, the resulting statistical description is equivalent to the one of a specific Ising model with frustrated spins [8, 9]. Therefore, the MEM approach can be interpreted as an effective mapping of the dynamics from an out-of-equilibrium, multi-component system into a “Hamiltonian” one [11], specified by relatively few parameters when compared to the full set of possible states of the system in question, allowing to apply the framework of thermodynamics. Within the scope of neuroscience, MEM has been mostly used to analyse experimental data of neuronal recordings, such as visual inputs from cells of a salamander [8, 11, 12, 18–20] and rat retina [21], numerical simulations from a phenomenological model of retinal ganglion cells [22], responses from hippocampus place cells in rodent brains [13, 15], synchronized and desynchronized neuronal activity from the primary visual cortex of anaesthetised cats and awake monkeys [23], *in vivo* and *in vitro* neuronal activity from cortical tissue of rodents [24] and the nervous system of the nematod *C. Elegans* [14], an organism whose pattern of connectivity between all its 302 neurons is well known [25]. Overall, these studies have given insights regarding collective [8] and functional characteristics [19] of biological neural networks, as well as helping unraveling the information content of neuronal responses [11, 12, 26] and its analogous thermodynamic properties [12]. Particularly, it was shown that the thermal fluctuations present in these Ising systems tend to diverge with system size [9, 12, 14, 23, 24], a sign of critical behaviour. However, experimental measurements in real networks can pose challenges that may impact the relevance and range of applicability of the conclusions drawn using the MEM method. For instance, contemporary neuronal recording techniques are restricted by the duration and sampling rate of the recordings [20, 27] and a precise association between which neuron generated which spike is an open problem, known as “spike sorting” [28], possibly affecting the accuracy of the measured correlation functions [29]. Furthermore, precise estimations of the fraction of excitatory and inhibitory neuronal populations in real systems are also often difficult to obtain [30]. To mitigate these challenges, numerical models can offer a more controlled environment for studying neuronal activity, allowing one to systematically change many parameters of choice, tuning the activity state, make systematic size studies which do not rely on subsampling, and generate many equivalent independent samples, thus producing statistically relevant data.

In this context, we apply the MEM method to describe spontaneous neuronal activity generated by an Integrate-and-fire (IF) model implemented on a scale-free network with short- and long-term plasticity [31]. The IF model relies on a tuning parameter that regulates the dynamical state of the system [31, 32]. At appropriate values of the tuning parameter, the model generates bursts of firing activity with characteristic spatio-temporal statistics, known as neuronal avalanches, first observed in 2003, in acute slices of rat cortex [33], displaying sizes *S* and durations *D* that are power-law distributed, as *P* (*S*) *∝S*^*−*1.5^ and *P* (*D*) *∝ D*^*−*2^, suggesting that biological neural networks operate near a critical point, as proposed in previous numerical and analytical studies [34–36], and is consistent with the divergence of thermal energy fluctuations observed in relatively small Ising-like models constructed from experimental neuronal data using the MEM method [9, 12, 14, 23]. Results suggest that the brain might act between a quiescence-like state and a state of hyperactivity, possibly offering several biological advantages such as optimal information transmission and storage [37]. Signs of criticality have been observed in a wide variety of animals, including humans [38–41], monkeys [41, 42], cats [23, 43], salamanders [44], turtles [45], worms [14, 46] and fish [47], a hallmark of universality [34].

Our aim here is to construct, using the MEM method mentioned above, pairwise, fully-connected Ising models of frustrated spins with local fields that replicate the average local activities and correlation functions between the neurons of the IF model in the critical state. Then we investigate spontaneous neuronal activity under a thermodynamic framework using our biologically relevant numerical IF model. In particular, we will investigate how the parameters of the Ising model and its thermodynamics properties change with the size and connectivity of the neuronal network. Additionally, we examine how different fractions of inhibitory neurons may influence the associated Ising models. We also consider partially-connected spin networks with only a subset of the total pairwise interactions, associated with the strongest neuronal pairwise correlations, to be able to analyse networks of sizes larger than the ones typically considered so far in the literature [27].

The following section is organized in 6 subsections. The first one analyses the firing dynamics generated by the IF model. The subsequent ones present the results from the mapping of the IF model into an Ising-like model using the MEM method and a study of its associated properties. In the penultimate section, we present a general discussion of the results and conclusion. In the final section, we describe the details of the IF model implementation and the MEM method.

## Results

We consider scale-free neural networks of different size *N∈* [20, 500] with short- and long-term plasticity, as well as a refractory mechanism, where neurons remain inactive for a single timestep immediately after firing (see Methods). Together with plastic adaptation, the refractory mechanism has been shown to play a crucial role in the appearance of critical behaviour and in shaping the topology of the network [48]. All measurements are performed on networks in the critical state. This is done by appropriately tuning the neurotransmitter recovery parameter *δu*_rec_ (see “Integrate-and-fire model” in Methods) and verifying that the distributions of neuronal avalanche sizes 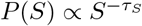 and durations 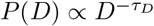 decay as power laws with exponents *τ*_*S*_ = 1.5 and *τ*_*D*_ = 2 (see S1 Fig and S2 Fig), characteristic of the mean field self-organized branching process [49], and consistent with both numerical and experimental observations [31, 33, 40, 50]. Initially, we consider only fully-excitatory networks. Later, we will also consider networks with a fraction *p*_in_ *>* 0 of inhibitory neurons. The time evolution of a system with *N* = 120 neurons in the critical state is shown in the raster plot of Fig 1.

**Fig 1.**
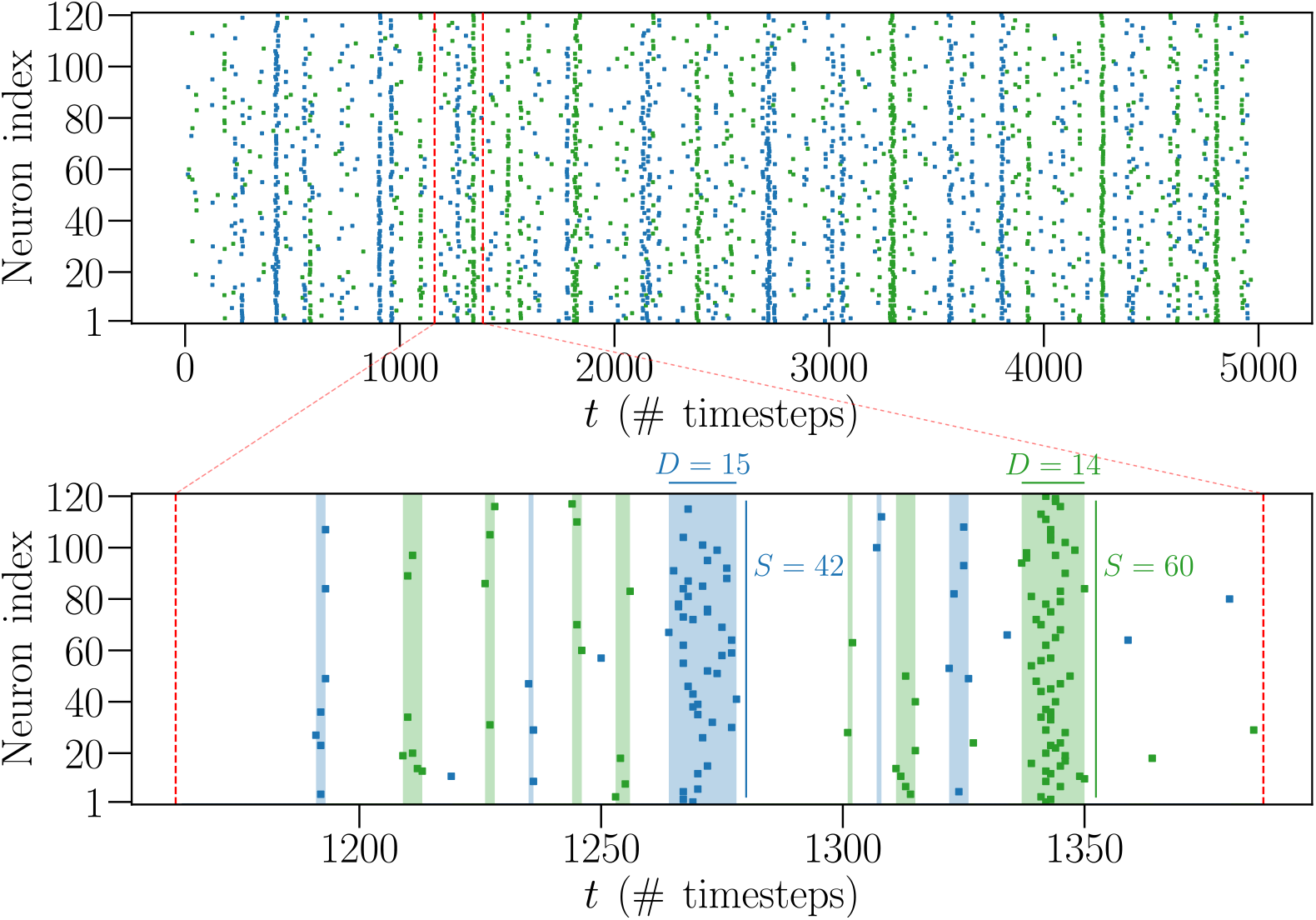
Time series for a system at criticality with *N* = 120 neurons. The top raster plot presents the time evolution of the firing states of *N* = 120 neurons over a certain number of timesteps, from a fully-excitatory IF network in the critical state. A coloured dot indicates that the respective neuron fired during that timestep, while an absence of a dot means it was inactive. The alternating green and blue colours are just a visual aid to distinguish between different avalanches. The bottom raster plot is a zoom-in of the top one, where the coloured shaded areas indicate avalanches of size *S >* 1 and duration *D*.

First, we analyse the firing statistics of the IF model networks. Then, we apply the MEM method to map the dynamics of the IF model networks of size *N∈* [20, 120] into a pairwise Ising model, defined by *N* “local fields” *h*_*i*_ and *N ·* (*N −* 1)*/*2 “interaction constants” *J*_*ij*_, and study its thermodynamical properties. More specifically, this mapping is achieved by using the so-called Boltzmann Machine (BM) algorithm. This learning algorithm searches for the set of parameters *h*_*i*_ and *J*_*ij*_ that best fit the pairwise Ising model to the data of the IF model by comparison with the average local activities ⟨*σ*_*i*_ ⟩ and correlation functions *C*_*ij*_ generated by each model (see “Maximum Entropy Modelling” in Methods).

### Firing statistics

To measure the IF network firing statistics, we consider a time bin Δ*t*_*b*_ = 5 timesteps. We assign a binary value 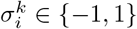 to each neuron at each time bin according to the following rule: 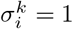 if neuron *i* fired at least once during the *k*-th time bin, 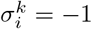 if the neuron *i* was inactive during the whole time bin *k*. We then calculate the average local activity ⟨*σ*_*i*_⟩ of the *N* neurons over *N*_*b*_ time bins, defined as

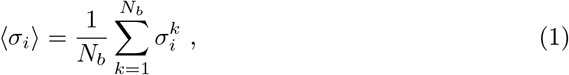

as well as the average two-point activity ⟨*σ*_*i*_*σ*_*j*_⟩ between all *N ·* (*N −* 1)*/*2 pairs of neurons *i* and *j*

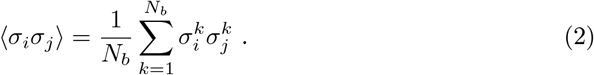

From these quantities we can also define the two-point correlation functions *C*_*ij*_,

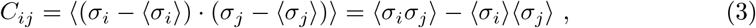

where ⟨*σ*_*i*_⟩ is closely related to the firing rate *r*_*i*_ = (⟨*σ*_*i*_ ⟩ + 1) */*2Δ*t*_*b*_ of neuron *i* [11], whereas *C*_*ij*_ quantifies the tendency for neurons *i* and *j* to fire together in the time interval Δ*t*_*b*_. Notice that, although likely related, the *C*_*ij*_ are completely distinct from the synaptic strengths *w*_*ij*_. For instance, the *C*_*ij*_ are defined for each pair *i* and *j* and are symmetric by definition (*C*_*ij*_ = *C*_*ji*_), whereas *w*_*ij*_ are not. For all simulations we use *N*_*b*_ = 10^7^.

In Fig 2 we show the distributions of ⟨*σ*_*i*_⟩ and *C*_*ij*_ obtained from *N*_*c*_ different network configurations (see S1 Fig for the respective avalanche statistics), for systems with different number of neurons *N*. We vary *N*_*c*_ according to *N* so that the total number of neurons considered *NN*_*c*_ *∼* 10000. The average local activity ⟨*σ*_*i*_⟩ is negative for all neurons (Fig 2a-c), implying that firing is a relatively rare event when considering individual neurons. In Fig 2d-f, we show that the correlation functions are small, but mostly non-zero for all neuron pairs, with their distributions peaked near zero, becoming sharper as the system size *N* increases. It is also noteworthy that, in experimental studies of the vertebrate retina [8], even weak pairwise correlations have been shown to likely become statistically significant in large networks, influencing the dynamics at the scale of the whole network. Another useful quantity that characterizes the firing dynamics is the probability *P* (*K*) to observe *K* ∈ [0, *N*] neurons firing simultaneously within a time bin Δ*t*_*b*_,

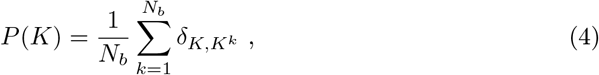

where 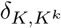 is the delta Kronecker function and 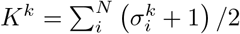 counts the number of neurons that fired during the *k*-th time bin. In Fig 3 we plot the probability (4) for several networks of different size *N*. For all *N, P* (*K*) is well fitted by an exponential distribution in an intermediate regime. The exponential factor of the distributions consistently decreases as *N* increases, indicating that it is more likely to observe concurrent firing between neurons in larger networks.

**Fig 2.**
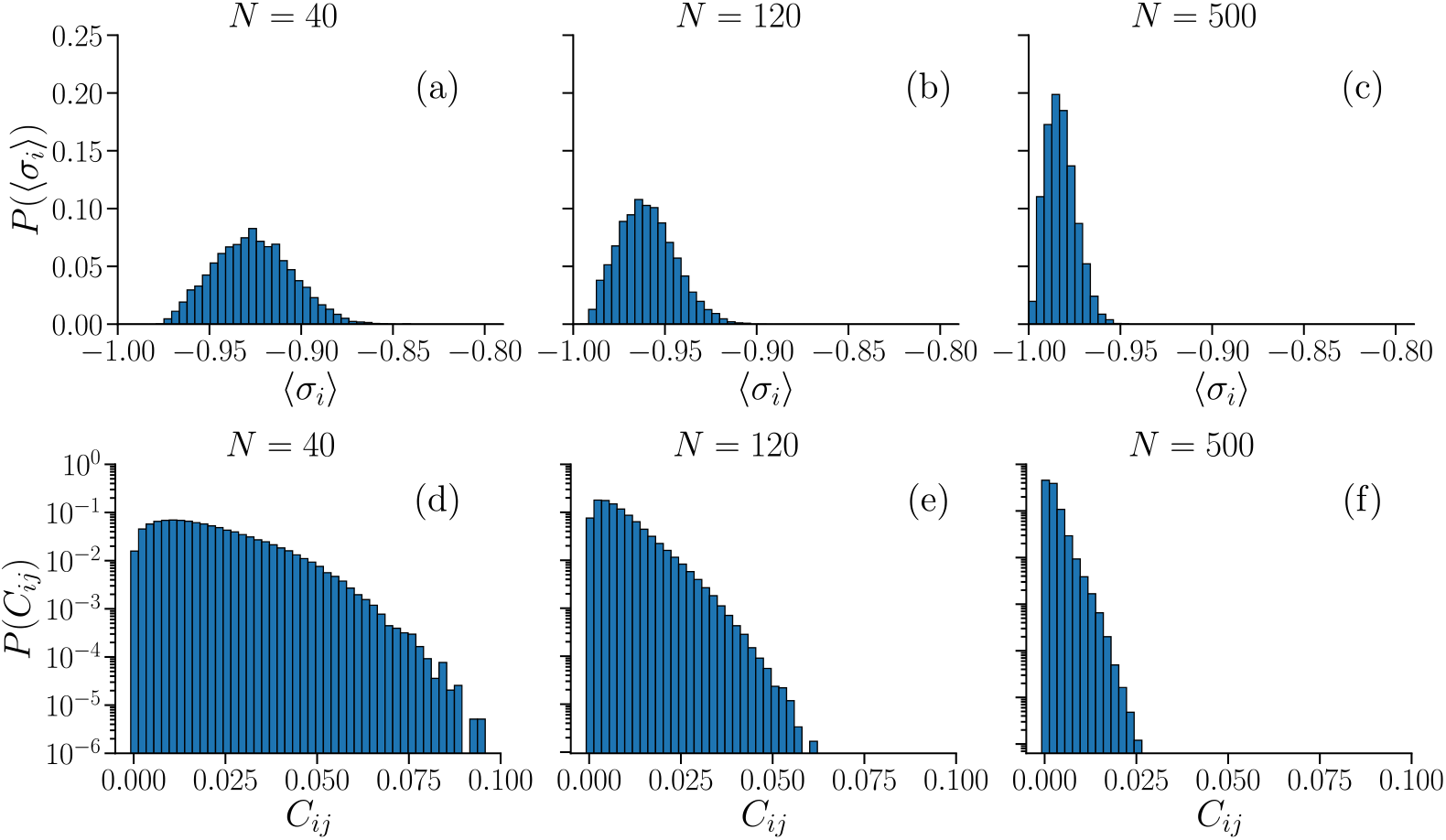
Distributions of the average local activity ⟨*σ*_*i*_⟩ and of the correlation functions *C*_*ij*_ in IF networks. Average local activity ⟨*σ*_*i*_⟩ (a-c) and correlation functions *C*_*ij*_ (d-f), calculated as averages over *N*_*b*_ = 10^7^ time bins. Distributions are obtained for sizes *N* = {40, 120, 500}, for *N*_*c*_ *∼* 10000*/N* different fully-excitatory networks.

**Fig 3.**
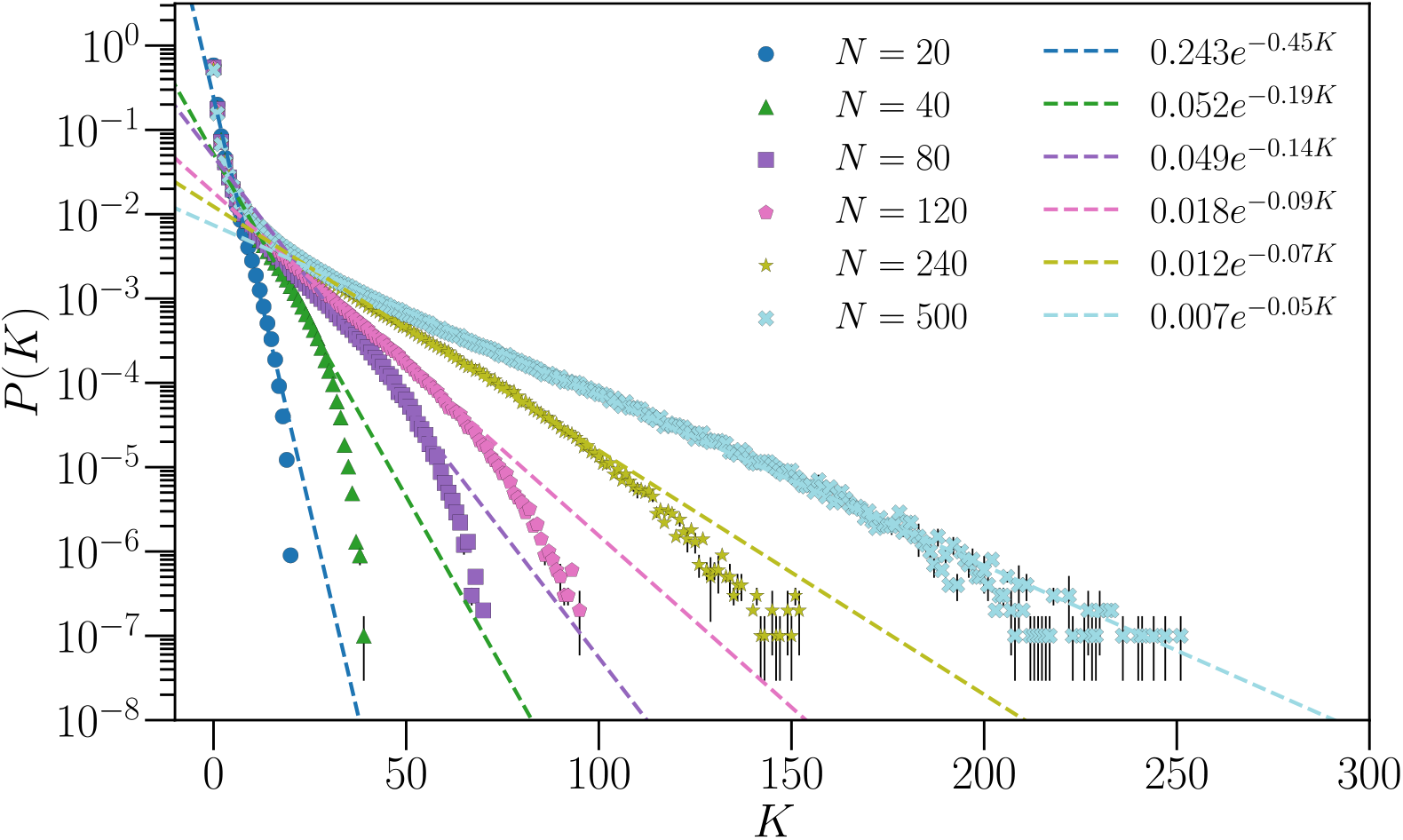
Probability *P* (*K*) of observing *K* neurons firing simultaneously during a time bin Δ*t*_*b*_ = 5 timesteps. Results are averages over *N*_*c*_ different fully-excitatory IF networks configurations of size *N*. *N*_*c*_ varies with *N* as in Fig 2. For each configuration, *P* (*K*) is estimated by averaging over *N*_*b*_ = 10^7^ bins. Error bars are given by the standard error, and most of them are smaller than the symbol size.

### Results of the Boltzmann Machine learning

Next, we consider pairwise Ising models of different system sizes *N* = {20, 40, 80, 120}, fitted to reproduce the temporal averages generated by the IF model, using the Boltzmann Machine (BM) algorithm (see “Maximum Entropy Modelling” in Methods). Each spin *i* can either be in the up-state (*σ*_*i*_ = 1) or down-state (*σ*_*i*_ = *−* 1), defining a particular spin configuration ***σ*** = {*σ*_1_, *σ*_2_, …, *σ*_*N*_ }. The Ising Hamiltonian *H*(***σ***) is given by

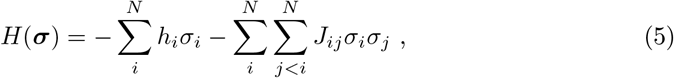

with *h*_*i*_ and *J*_*ij*_ the fitting parameters, acting as local fields on spin *i* and interaction constants between spins *i* and *j*, respectively. When appropriate, we use the superscript (IF) and (BM) to distinguish between measurements using the IF model and those obtained with the BM using Monte Carlo. More specifically, ⟨ … ⟩ ^(IF)^ indicates an average over time bins in the IF model while ⟨… ⟩^(BM)^ is an average over spin configurations of the Ising model.

In Fig 4 we present the values of {*h*_*i*_} and the distributions of the interaction constants {*J*_*ij*_ }, fitted to reproduce the temporal averages {⟨*σ*_*i*_⟩ ^(IF)^ } and {⟨*σ*_*i*_*σ*_*j*_ ⟩ ^(IF)^ } of fully-excitatory IF networks of size *N*, tuned to the critical state. More specifically, Figs 4a-d show ranked plots of the fields *h*_*i*_, where *h*_1_ (*h*_*N*_) corresponds to the spin associated with the neuron that fired the most (least) in the IF model. The fields *h*_*i*_ are mostly negative for all *N*. This is not surprising considering that a neuron in the IF model fires rarely, as discussed in previous sections, and the negative fields are a consequence of this sparse activity. We note, however, that, as the system size *N* increases, an increasingly larger fraction of spins have positive *h*_*i*_, corresponding to the spins associated with the most active neurons. In Figs 4e-h we plot the distributions of the learned couplings *J*_*ij*_. The distributions are bimodal, with the largest peak near zero and the second one at positive *J*_*ij*_. However, the height of the second peak appears to decrease with the system size *N*, and the limiting distribution when *N→∞* becomes consistent with a normal distribution centered around zero. It is interesting to recall that random, normally distributed couplings *J*_*ij*_ characterize the so-called Sherrington-Kirkpatrick (SK) model [51, 52], an Ising-like model which has been shown to exhibit both ferromagnetic and spin glass phases [51]. Unlike the SK model, however, the learned Ising models exhibit high heterogeneity in the distribution of the fields *h*_*i*_, and since the learning technique fits simultaneously both the fields and interaction constants, the inference of the {*J*_*ij*_} should not be considered numerically decoupled from that of the {*h*_*i*_}.

**Fig 4.**
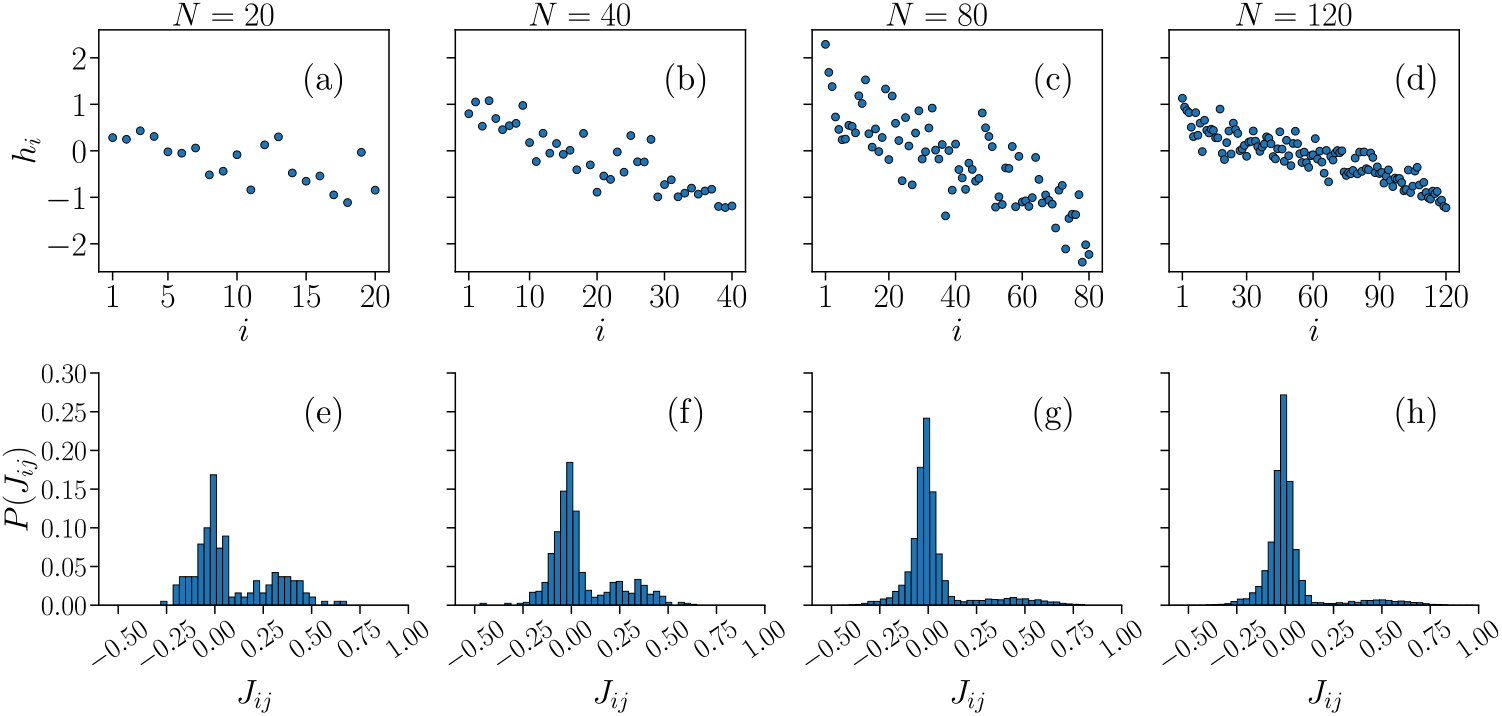
Sets of learned fields *h*_*i*_ and interaction constants *J*_*ij*_ of the pairwise Ising model that reproduce the time averages of the IF model. Plots of the fields *h*_*i*_ (a-d), sorted by the average local activity ⟨*σ*_*i*_ ⟩ ^(IF)^ of the associated neuron *i*, in order of decreasing ⟨*σ*_*i*_⟩ ^(IF)^, and distributions of the interaction constants *J*_*ij*_ (e-h), learned by the BM to reproduce the average local activity ⟨*σ*_*i*_ ⟩ and two-point correlation functions *C*_*ij*_ of a fully-excitatory IF neural network at criticality, with *N* = {20, 40, 80, 120} neurons.

To test the quality of the learning process, in Fig 5 we compare the {⟨*σ*_*i*_⟩ ^(BM)^} and 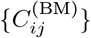 of the Ising-like model with the neural network data of the IF model {⟨*σ*_*i*_⟩^(IF)^} and 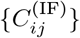. As described above (by virtue of equations (36) and (37), in the “Methods” section), if the learning is successful, these quantities should be identical since these are precisely the constraints imposed in the learning procedure. This is confirmed for both ⟨*σ*_*i*_ ⟩ and *C*_*ij*_, for all *N*, with all points following closely the bisector *y* = *x*.

**Fig 5.**
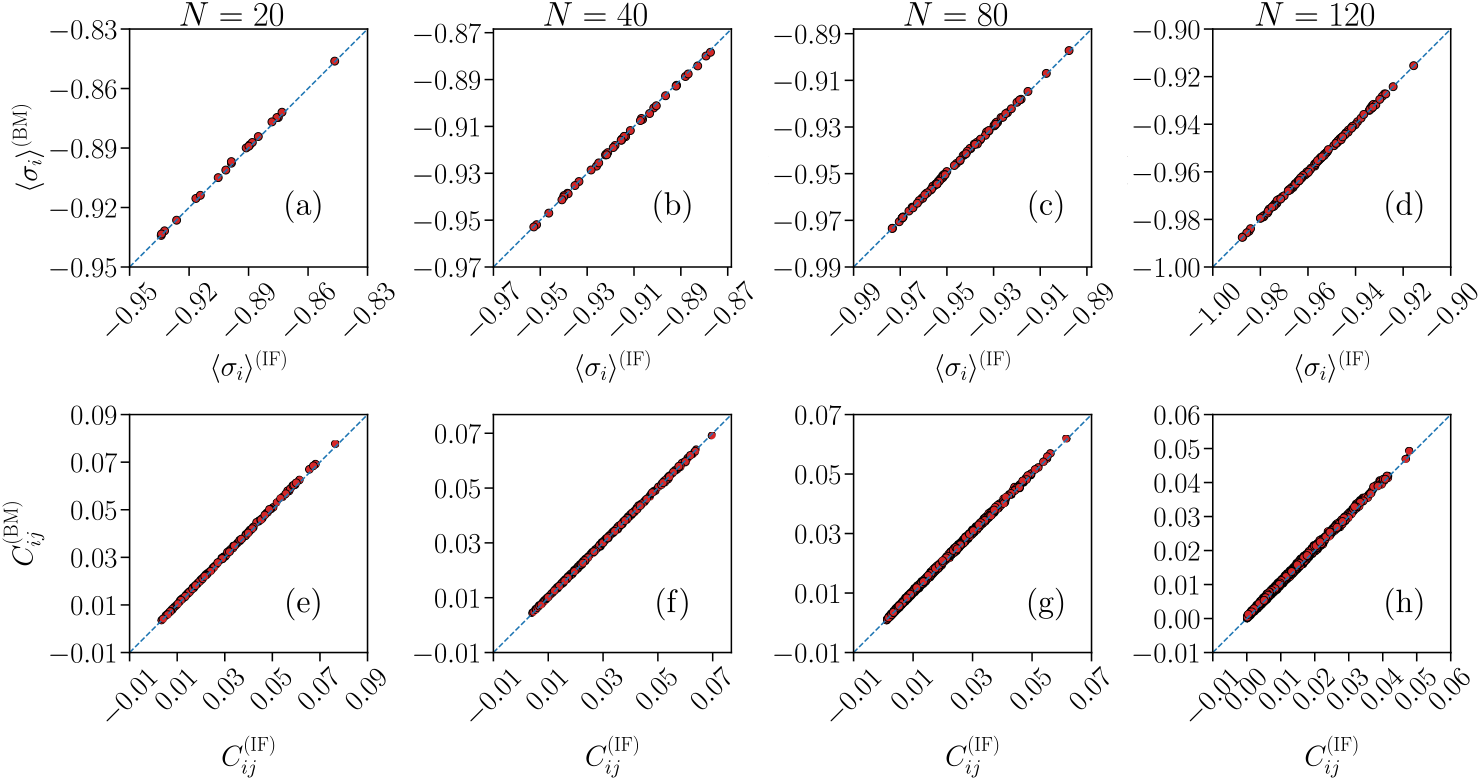
Quality test for the BM learning process. Comparison between the average local activity ⟨*σ*_*i*_ ⟩ (a-d) and correlation functions *C*_*ij*_ (e-h) of the Ising-like model (*y*-axes) and IF model network (*x*-axes), for system sizes *N* = {20, 40, 80, 120}. The blue dashed lines are the bisector *y* = *x*. Results are averages over *N*_*b*_ = 10^7^ time bins for the IF model (IF), and averages over *M*_*c*_ = 3 *·*10^6^ spin configurations for the Ising model (BM). Error bars are given by the standard error, and are smaller or equal to the symbol size.

An interesting quantity to assess the predictive power of the model are the three-point correlation functions *T*_*ijk*_ between all *N·* (*N −* 1) *·* (*N −* 2)*/*6 triplets of neurons *i, j* and *k*,

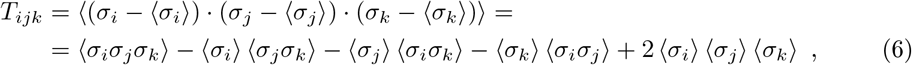

where the averages ⟨ … ⟩ are defined analogously as in equations (1) or (3). In Fig 6 we compare the triplets between the IF and the Ising models, for systems with *N* = {20, 40, 80, 120} considered previously. Even though the *T*_*ijk*_ are not constrained by the learning procedure, the generalized Ising model still captures their systematic qualitative behaviour. Unlike ⟨*σ*_*i*_ ⟩ and *C*_*ij*_, however, the triplets appear to be consistently overestimated by the Ising model, when compared to their IF model counterpart. We note that this behaviour was also seen in experimental recordings from stimulated neurons in a salamander retina [9], albeit to a less significant degree.

**Fig 6.**
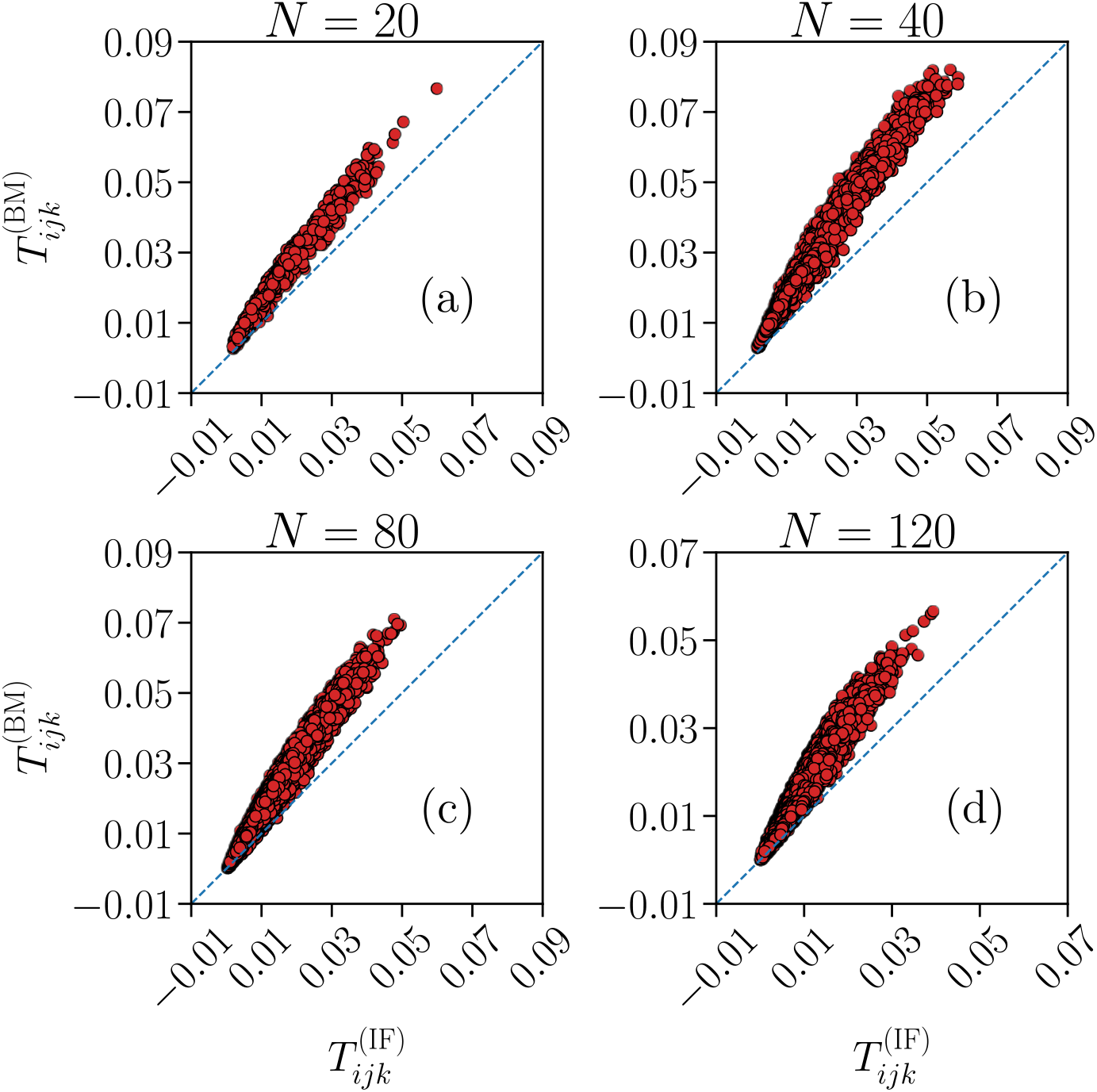
Predictive capability of the Ising model for the three-point correlation functions *T*_*ijk*_. Comparison between the three-point correlation functions *T*_*ijk*_ of the Ising-like model (y-axes) and the IF model network (x-axes), for system sizes *N* = {20, 40, 80, 120}. The blue dashed lines are the bisector *y* = *x*. Results are averages over *N*_*b*_ = 10^7^ time bins for the IF model (IF), and averages over *M*_*c*_ = 3 *·* 10^6^ spin configurations for the Ising model (BM). Error bars are given by the standard error, and are smaller or equal to the symbol size.

Another useful measure to assess how well the Ising-like model describes the IF model data is the probability for simultaneous firing *P* (*K*) within a small time window, as defined in equation (4). In the Ising model, the sum over time bins in equation (4) is replaced by a sum over spin configurations generated by Monte Carlo simulation, where *K* now counts the number of up-spins in a single configuration in the Monte Carlo evolution. We compare this quantity between the IF and Ising models in Fig 7 for systems of size *N* = {20, 40, 80, 120}. The Ising model seems to predict the *P* (*K*) correctly only when *K* is very small, of the order *K/N* ≲ 0.10, with noticeable discrepancies seen also for this range of *K*, particularly for the probability of no activity (*K* = 0), in systems of size *N >* 20. For larger *K*, there is an intermediate range where the Ising model underestimates *P* (*K*), followed by a region in which *P* (*K*) is greatly overestimated. This effect seems to increase with the system size *N*. This particular behaviour was also observed when training a pairwise Ising model to reproduce the time averages of stimulated neuronal activity from a salamander retina [11].

**Fig 7.**
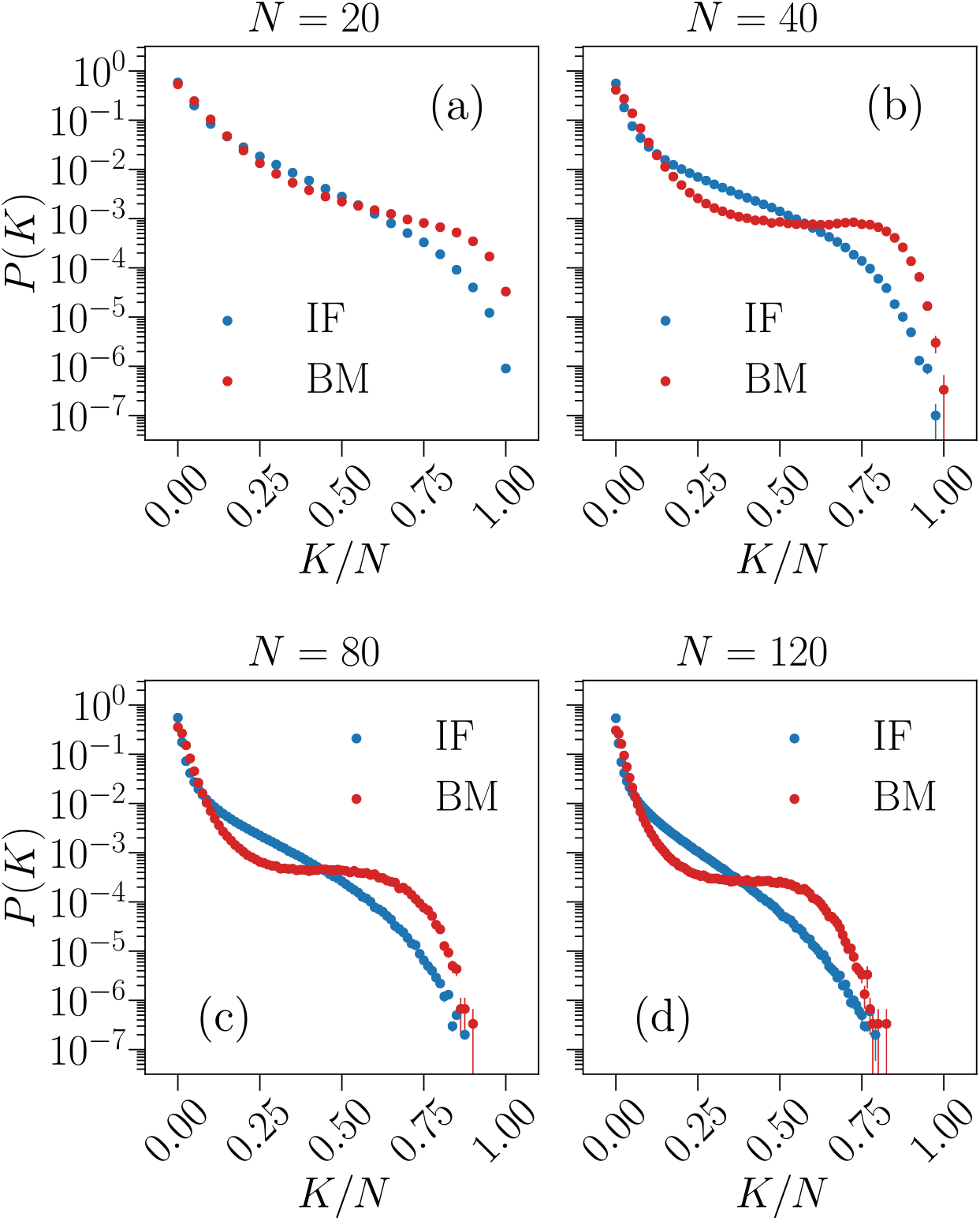
Predictive capability of the Ising model for the probability of simultaneous firing *P* (*K*). Comparison between the simultaneous firing/up-state probability *P* (*K*) of the Ising-like model (red) and the IF model network (blue), for system sizes *N* = {20, 40, 80, 120 }. Results are averages over *N*_*b*_ = 10^7^ time bins for the IF model (IF), and averages over *M*_*c*_ = 3 *·*10^6^ spin configurations for the Ising model (BM). Error bars are given by the standard error, and most are smaller or equal to the symbol size.

### Thermodynamics of the pairwise Ising models

Having mapped the neural network to a pairwise Ising model, we can start to analyse its properties. Given a system with *N* spins, for each spin configuration we can measure the magnetization 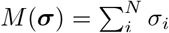 and the energy 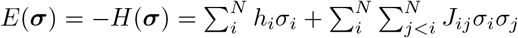. From the fluctuations of these quantities, according to the fluctuation-dissipation theorem, we can calculate the susceptibility *χ* and the specific heat *C*_*v*_ of the Ising system as a function of temperature *T*,

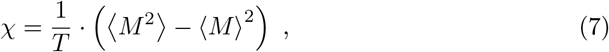

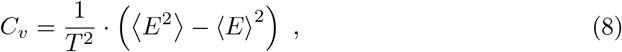

where ⟨…⟩ indicates an average over spin configurations generated by Monte Carlo simulations. By changing *T*, one can probe the thermodynamic properties of these spin systems. We start from a random spin configuration and thermalize it as described in the Methods section. Notice, however, that the MEM method only claims that the Ising model is representative of the IF model for *T* = 1 *≡ T*_0_, since that is the temperature used during the BM learning procedure. Since there is no evident analogy between the control parameter of the learned Ising models, *T*, and that of the IF model, *δu*_rec_, temperatures *T* ≠ *T*_0_ have no obvious physical meaning in these learned Ising models, besides being a useful parameter to check if there is anything special about this *T*_0_ [12].

In Fig 8 we plot the average magnetization per spin *m* = ⟨*M* ⟩ */N*, the susceptibility *χ* and the specific heat *C*_*v*_ as a function of the temperature *T* for systems of size *N* = {20, 40, 80, 120}. For each *N*, we consider five Ising systems with distinct parameters learned from different IF networks of identical size at criticality, to assess the sample-to-sample variations for IF networks with different specific connectivities between the neurons.

**Fig 8.**
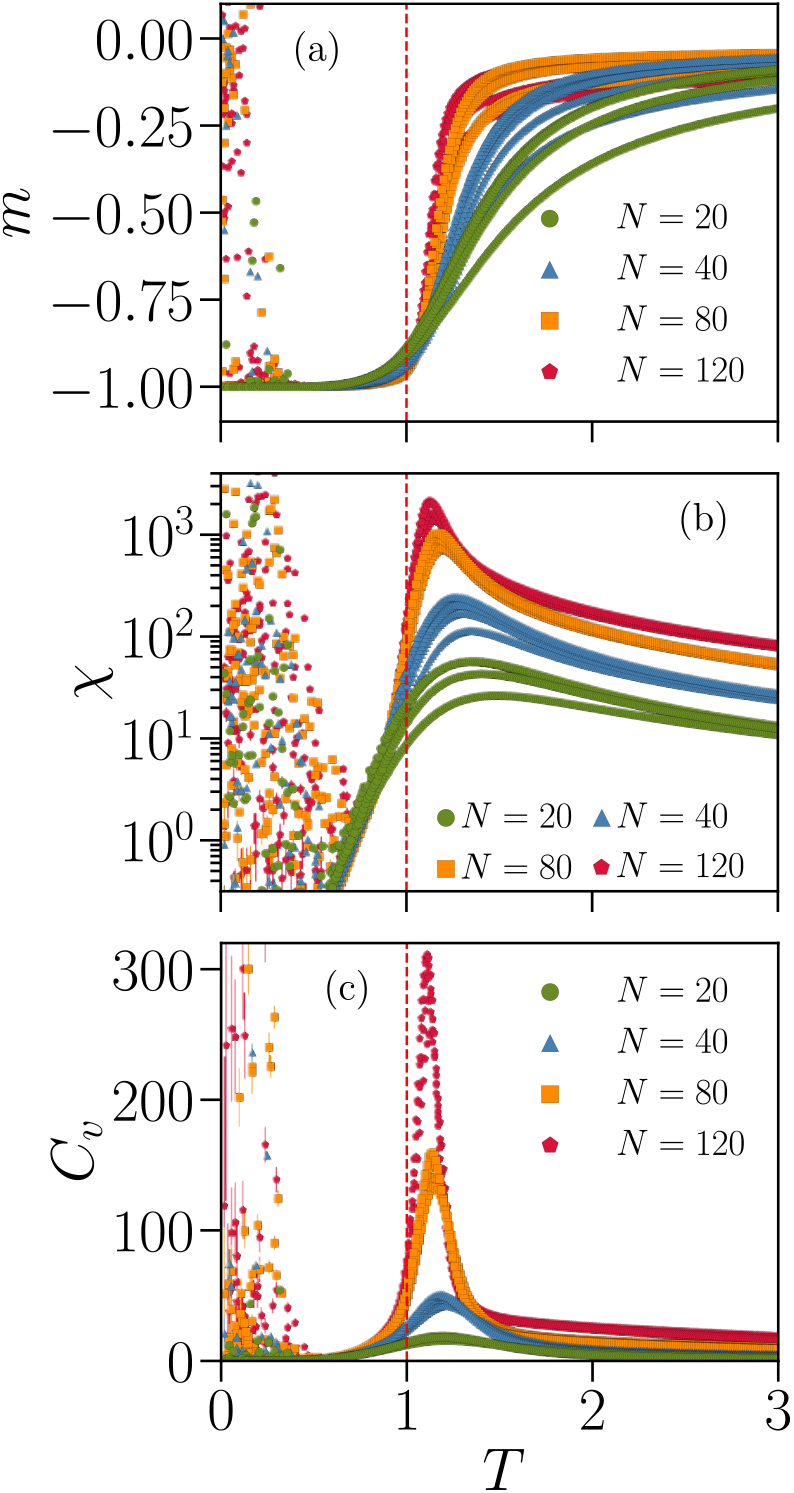
Thermodynamic functions of Ising-like models associated with fully-excitatory IF networks of different system sizes *N* . Average magnetization per spin *m* (a), susceptibility *χ* (b) and specific heat *C*_*v*_ (c) as a function of the temperature *T ∈* [0.1, 3.0], for systems with *N* = {20, 40, 80, 120} spins. Different curves with the same colour and symbol correspond to Ising systems with parameters fitted to different IF network configurations with the same size *N* but different specific connectivities. The vertical dashed lines indicate *T* = *T*_0_ = 1, the “default” temperature used in the BM learning procedure to fit the respective Ising parameters to each IF network. The cloud of random values observed for *T <* 1 suggests the presence of a spin-glass phase, where the thermal energy is insufficient to drive the system away from the initial random spin configuration. Results are averages over *M*_*c*_ = 3 *·*10^6^ spin configurations. Error bars are given by the standard error and are overall smaller or equal to the symbol size.

The maxima of the susceptibility and specific heat appear near the “default” temperature *T*_0_ = 1 associated with the IF model neuronal data, and its height increases with *N*, indicating that the neural networks tend to maximize thermodynamic fluctuations. This phenomenon, where fluctuations diverge with increasing system size, is a hallmark of a system operating near a critical point [53].

Looking now at different network configurations of the same size *N*, we can clearly detect large variations across configurations. Particularly, for different IF model networks with the same *N*, the maximum value of the susceptibility *χ* varies by a factor ∼2 for sizes *N* = {20, 40 } and ∼1.5 for *N* = {80, 120}.

Another interesting result is the cloud of random values observed for *T <* 1 for all thermodynamic quantities. This may be an indication that, for low temperatures, the Ising model transitions into a spin-glass phase. Since this phase usually exhibits complex energy landscapes [52], thermal fluctuations in this temperature regime might not be sufficient to drive the system away from the randomly chosen initial configuration and into the ground state. Indeed, starting with all *σ*_*i*_ = *−* 1 as initial configuration for the Monte Carlo simulations removes this effect entirely (see S3 Fig).

### Subnetworks

In experiments, neuronal recordings are usually performed over a subset of neurons out of the total population [11, 15]. As such, it is of practical interest to study how network subsampling might affect the analysis.

A very recent study [15] shows that the properties of firing statistics change with the spatial distribution of the subsampled neuronal patch. Taking this into account, we will consider two types of subnetworks, also studied in the aforementioned work: a subset of neurons that are spatially close together, Fig 9a, or picked at random positions, Fig 9b. For the former case, we select the neurons that are closest to the center of the cubic lattice.

**Fig 9.**
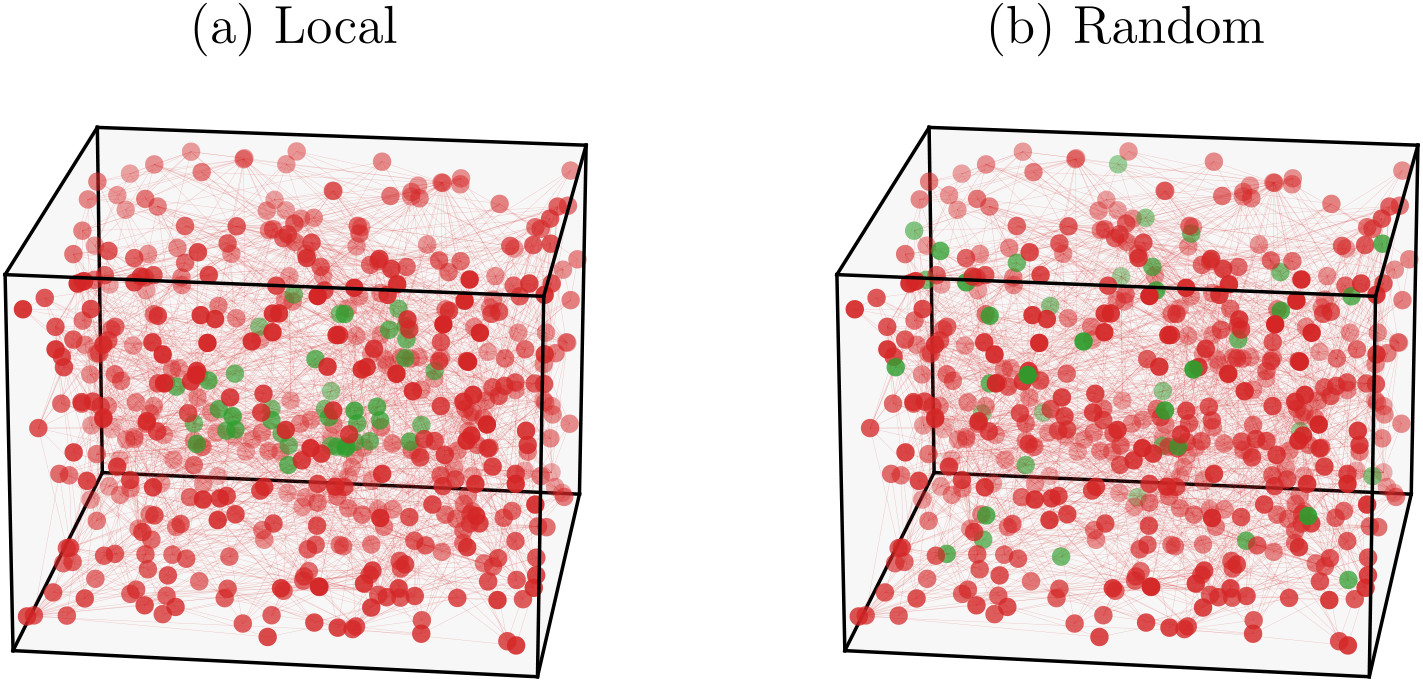
Schematic representations of IF model subnetworks with different spatial distribution. Subnetworks (green) with neurons closely packed together (a) or picked at random positions (b). Subnetworks have *n* = 40 neurons in a system of size *N* = 500.

In Fig 10 we plot the results of the simultaneous firing probability *P* (*K*) and triplets *T*_*ijk*_ measured in subnetworks of *n* = 40 neurons in systems with *N* = 500 total neurons, for both spatial distributions described previously. We also compare these quantities with the predictions of the corresponding Ising models. The learning efficiency for the parameters of these models is similar to the cases using full networks, as assessed by inspecting the constrained temporal averages, ⟨*σ*_*i*_⟩ and *C*_*ij*_ (see S4 Fig).

**Fig 10.**
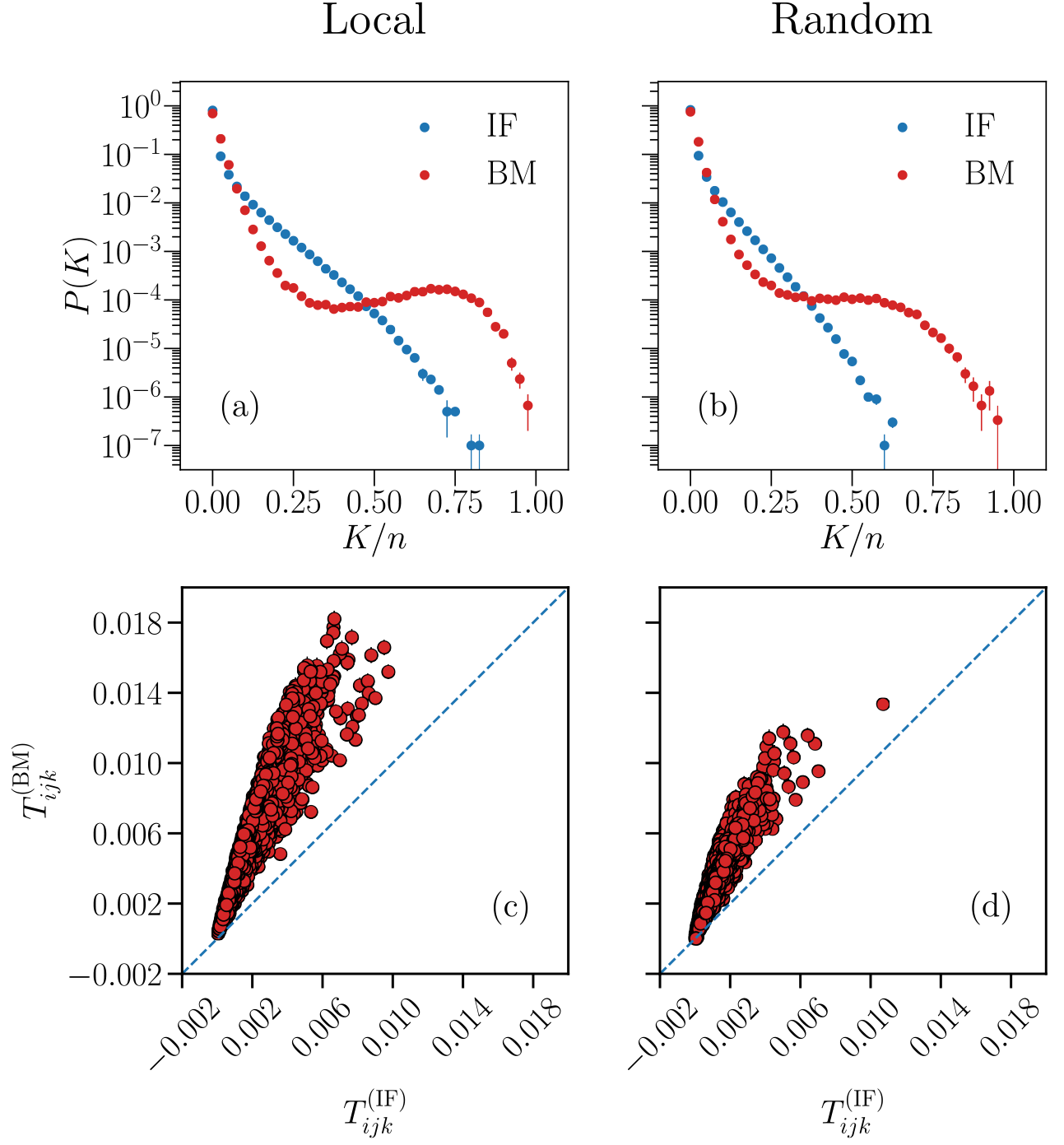
Predictive capability of the Ising model for IF model subnetworks. Comparison between Monte Carlo sampling of the Ising model (BM) and the neural network data (IF) of the probability *P* (*K*) (a-b) and triplets *T*_*ijk*_ (c-d), for subnetworks with *n* = 40 neurons, with a local (a and c) and random (b and d) spatial distribution, in a system with *N* = 500 neurons. The blue dashed lines in the bottom plots are the bisector *y* = *x*. Results are averages over *N*_*b*_ = 10^7^ time bins for the IF model (IF), and averages over *M*_*c*_ = 3 *·* 10^6^ spin configurations for the Ising model (BM). Error bars are given by the standard error, and are overall smaller or equal to the symbol size.

As in the case of entire networks, the Ising models fail to predict *P* (*K*) for large *K*, with even larger discrepancies (Fig 10). For the IF network, the *P* (*K*) distribution decays exponentially for both subnetworks, as for the entire network case. Analogously, the probability *P* (*K*) for the learned Ising models shows an initial exponential decay for low *K*, for both spatial distributions. Conversely, at *K/n ≈* 0.35, the behaviour of *P* (*K*) from the Ising model is slightly different between the two spatial distributions. For the local case, the probability for observing simultaneously up-spins reaches a second local maximum at *K/n ≈* 0.75. Notice, however, that this is a small effect, magnified by the logarithmic scale. On the other hand, for the random case, a plateau in *P* (*K*) of the Ising model is observed in the intermediate regime of *K*, followed by a subsequent decay. These differences between the IF and Ising model, compared to the full network case (Fig 7), could be due to the fact that we are neglecting influences from neurons that are not included in the subnetworks, and therefore not encoded in the Ising model parameters.

The behavior of the triplets is similar to the case of entire networks (Fig 6), wherein the triplets are systematically overestimated by the Ising model. We note also that overall the triplets have much smaller values, when compared to the ones found for the full network with *N* = 40. Similar results were found for both *P* (*K*) and the *T*_*ijk*_ for subnetworks of larger size *n* = 80 in the same IF network with *N* = 500 (not shown). The fact that both cases of spatial distributions yield Ising models with similar accuracy in reproducing the original IF model data is unexpected, as this is seemingly in contrast to recent experimental studies on mouse brains [15], where the generalized Ising model predicted better the results from subgroups of *∼* 100 neurons that were spatially clustered together. It should be noted, however, that the experimental study pertains to stimulated neuronal activity due to visual stimuli, whereas our model simulates spontaneous neuronal activity [31], in the absence of any external stimulation.

### Networks with inhibitory neurons

Inhibitory neurons hamper neuronal activity, consequently affecting the dynamics of the network [54, 55]. Increasing the percentage *p*_in_ of inhibitory neurons in a neural network moves the system into a subcritical regime [55]. A way to keep the system close to the critical state when *p*_in_ *>* 0 is by increasing the value of the tuning parameter *δu*_rec_. Here we consider systems of size *N* = 80 and *p*_in_ *>* 0, in the critical state (see S2 Fig), and study how the presence of inhibition might affect the properties of the associated Ising-like models. The agreement of the average local activities and correlation functions between the Ising model and the IF networks with inhibitory neurons is the same as for fully-excitatory ones (see S5 Fig).

In Fig 11 we show the distributions of fields *h*_*i*_ and interaction constants *J*_*ij*_ learned for IF networks with *p*_in_ = {0%, 10%, 20%} inhibitory neurons.

**Fig 11.**
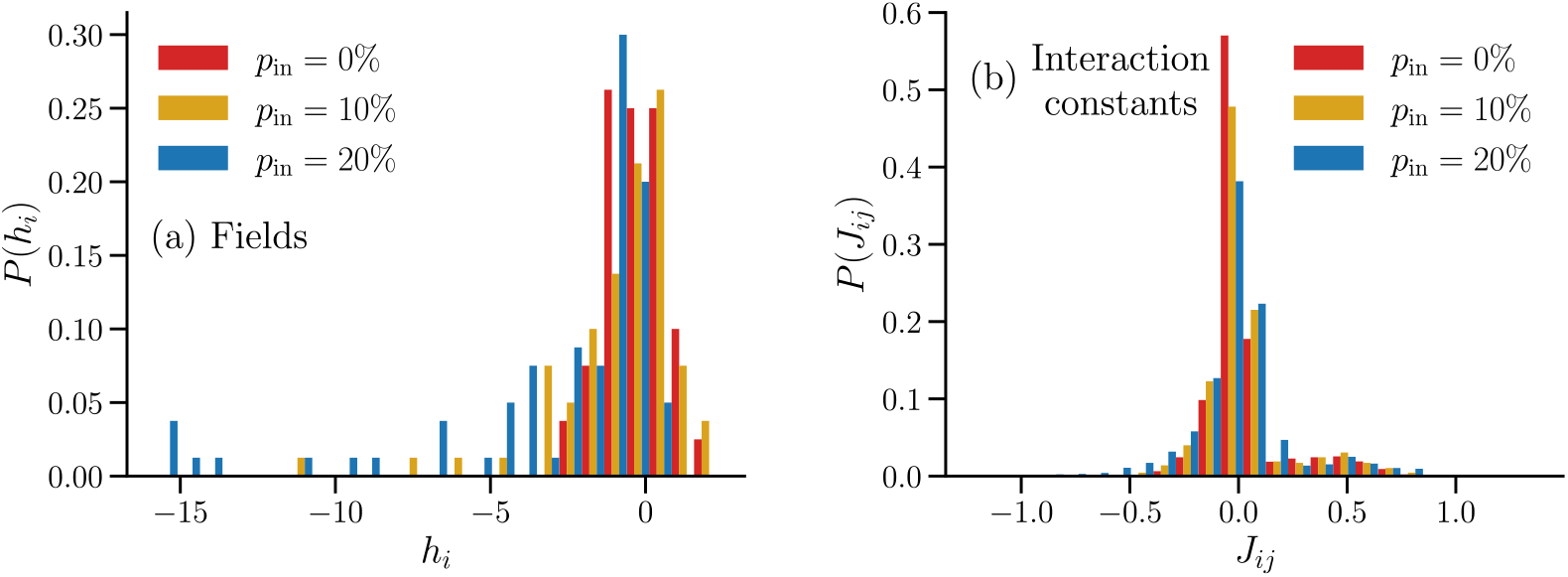
Distributions of the learned fields *h*_*i*_ and interaction constants *J*_*ij*_ of a pairwise Ising model for neural networks with different fractions of inhibitory neurons *p*_in_. Fields *h*_*i*_ (a) and interaction constants *J*_*ij*_ (b) associated with IF networks with *p*_in_ = {0%, 10%, 20% } and *N* = 80, obtained after *N*_BM_ = 60000 iterations of the BM.

As seen in Fig 11a, the distribution of fields *h*_*i*_ acquires a tail towards negative *h*_*i*_ for *p*_in_ *>* 0, becoming more pronounced as *p*_in_ increases. Interestingly, these *h*_*i*_ are associated with excitatory neurons (see S6 Fig). A possible explanation is that, in the IF network, the neurons with the smallest average local activity ⟨*σ*_*i*_ ⟩ are not necessarily inhibitory neurons, but rather neurons with incoming connections from inhibitory ones, which in turn are more likely to be excitatory since *p*_in_ *<* 50%. This indicates that the {*h*_*i*_} do not encode the information about whether a neuron is excitatory or inhibitory. In Fig 11b we see that the peak of the distributions of the interaction constants *J*_*ij*_, decreases with *p*_in_, with a more pronounced tail towards negative *J*_*ij*_ observed for *p*_in_ = 20%. Interestingly, the presence of the second peak of *P* (*J*_*ij*_) at positive *J*_*ij*_ seems to be robust with respect to changes in *p*_in_.

In Fig 12 we plot as a function of the temperature *T* the thermodynamic functions *m, χ* and *C*_*v*_ of the Ising systems with the parameters of Fig 11. In Fig 12a, the plateau for *m* at high *T* in the system associated to *p*_in_ = 20% is lower than the one associated to the fully-excitatory network, indicating a higher tendency for spins to be in the down state (*σ*_*i*_ = *−* 1) for the former case in the paramagnetic phase, which could stem from the inhibitory neurons. This is consistent with the observation of stronger negative fields *h*_*i*_ when *p*_in_ = 20%, as seen in Fig 11a. In Figs 12b and 12c, we see that the values of the maxima of *χ* and *C*_*v*_ decrease with *p*_in_, while their position with respect to the temperature remains unchanged. This suggests that introducing inhibition in the IF networks reduces thermal fluctuations in the associated Ising model.

**Fig 12.**
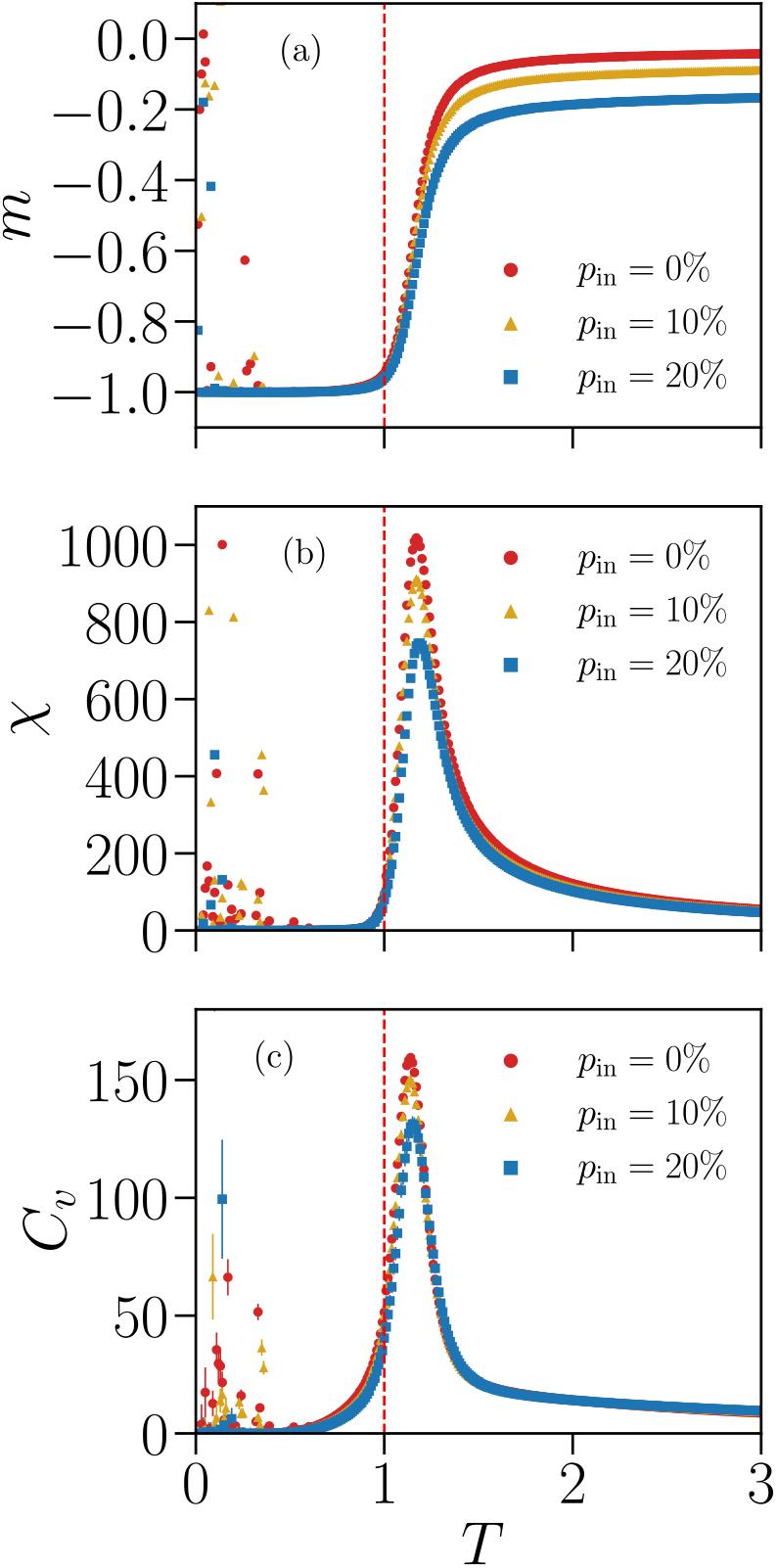
Thermodynamic functions of Ising-like models associated with IF model networks with different fractions of inhibitory neurons *p*_in_. Average magnetization per spin *m* (a), susceptibility *χ* (b) and specific heat *C*_*v*_ (c) as a function of the temperature *T* ∈ [0.1, 3.0], simulated using the learned parameters shown in Fig 11 for different fractions of inhibitory neurons *p*_in_ = {0%, 10%, 20% } and *N* = 80. As in Fig 8, the cloud of random values for *T <* 1 suggests the presence of a spin-glass phase. Results are averages over *M*_*c*_ = 3 *·*10^6^ spin configurations. Error bars are given by the standard error and are overall smaller or equal to the symbol size.

### Partially-connected pairwise Ising models

One of the main disadvantages of BM learning is its intense CPU time demand [27, 56], restricting the network sizes one could potentially analyse. To try to circumvent this problem, we will consider Ising models with pruned links [19], whereby we remove the couplings associated with the weakest correlated pairs of neurons. Specifically, we set *J*_*ij*_ = 0 if the corresponding 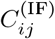 is below a certain threshold, defined as a fraction of the largest measured correlation function, i.e. if 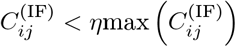, where *η ∈* [0, 1] sets the threshold. Thus, the BM only needs to learn a subset of the total number of couplings *J*_*ij*_, possibly accelerating the convergence process and consequently allowing the study of networks larger than the ones considered so far. Furthermore, an obvious speed-up is also achieved by making use of the fact that a subset of the {*J*_*ij*_ }are zero, and therefore can be disregarded in the double sum of equation (5) during the Monte Carlo simulations. We will consider three thresholds *η* ∈ {0.10, 0.15, 0.20 }, using a fully-excitatory system of size *N* = 180 in the critical state. Computing only a fraction of the interaction terms allowed to obtain the BM results for the *N* = 180 system in a less or comparable CPU time with respect to the one required by the *N* = 120 systems where we considered the full set {*J*_*ij*_}, with a greater CPU speed-up achieved the larger the threshold *η* is.

In Fig 13 we present the comparison between the correlation functions of the IF model and the partially-connected Ising model for the three different thresholds. We see that, even though more than 40% couplings *J*_*ij*_ have been removed in the Ising model for the lowest threshold considered, *η* = 0.10, and more than 70% for the largest one, *η* = 0.20, the overall data is still reconstructed by the partially-connected Ising model, with a decrease in quality of the fit for the lowest *C*_*ij*_ (grey dots in the plots) since they were not considered in the learning process of the BM. On the other hand, all ⟨*σ*_*i*_⟩ are well predicted by the Ising model (see S7 Fig) since we fit all the *N* = 180 fields {*h*_*i*_ }. In Fig 14 we plot the learned fields and the distributions of the learned non-zero interaction constants of the partially-connected Ising model for the three different thresholds *η*. While the set of fields {*h*_*i*_} remains qualitatively similar for the three thresholds *η*, with a majority of negative fields as in the case of the fully-connected Ising models, the distributions of the non-zero *J*_*ij*_ change significantly with *η*, with the peak at positive *J*_*ij*_ increasing and the one at *J*_*ij*_ *≈* 0 decreasing as we remove progressively more *J*_*ij*_. Since we remove only the *J*_*ij*_ associated with the smallest correlation functions *C*_*ij*_, this seemingly indicates that the second peak at *J*_*ij*_ *>* 0 also seen in the distributions for the fully-connected Ising models (Fig 4) is associated with the subset of the largest *C*_*ij*_.

**Fig 13.**
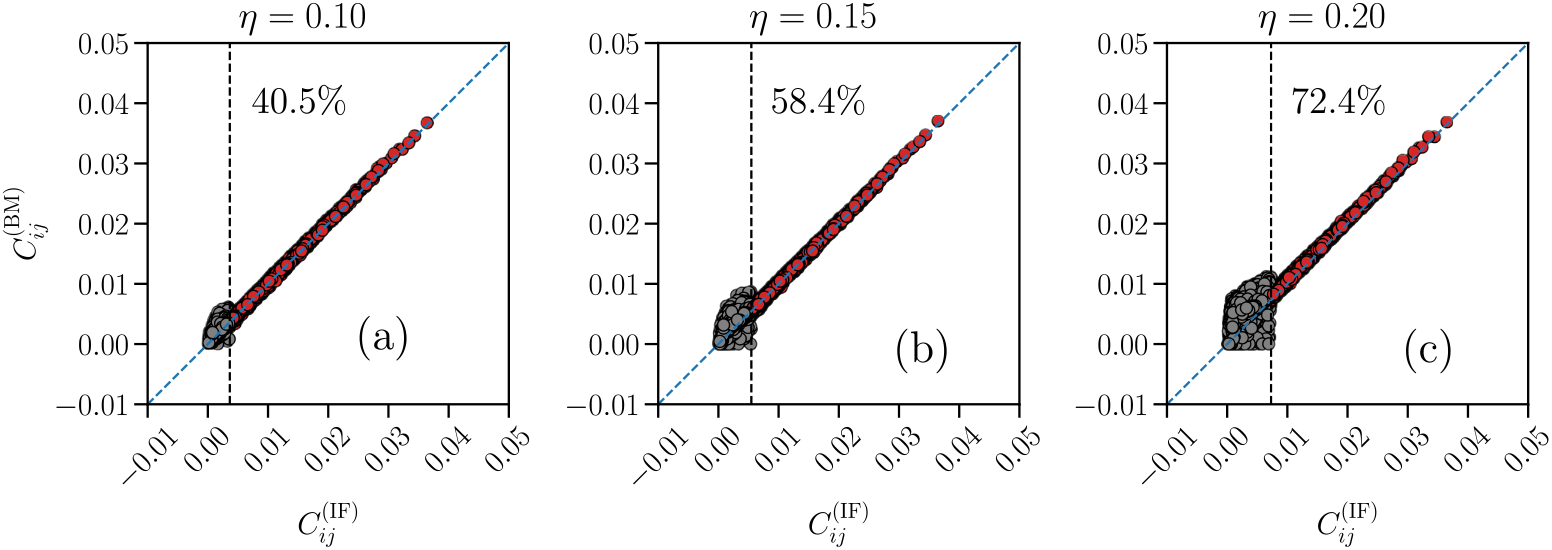
Quality test for the BM learning process with a partially-connected Ising model. Comparison between the correlation functions *C*_*ij*_ of the partially-connected Ising-like model (*y*-axes) and IF model network (*x*-axes), for a fully-excitatory system at criticality of size *N* = 180 and three different thresholds *η* = {0.10, 0.15, 0.20} for the removal of the *J*_*ij*_. If 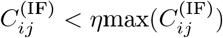 (grey dots), the corresponding interaction constant is set to *J*_*ij*_ = 0. The vertical dashed lines indicate the value 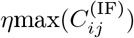, with the corresponding approximate percentage of removed *J*_*ij*_ reported at the right of this line. The blue dashed lines are the bisector *y* = *x*. Results are averages over *N*_*b*_ = 10^7^ time bins for the IF model (IF), and averages over *M*_*c*_ = 3 *·*10^7^ spin configurations for the Ising model (BM). Error bars are given by the standard error, and are smaller or equal to the symbol size.

**Fig 14.**
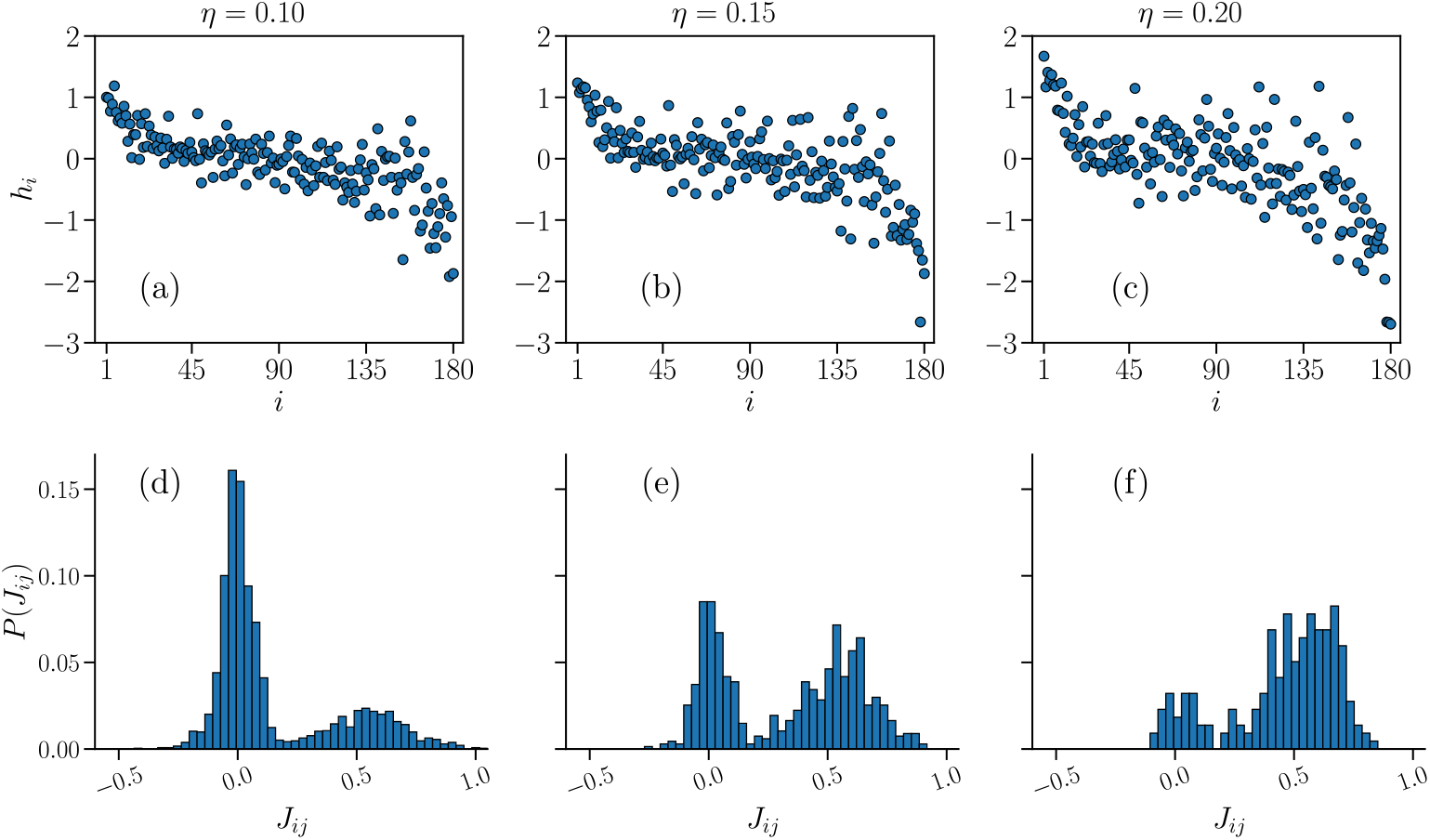
Sets of learned fields *h*_*i*_ and non-zero interaction constants *J*_*ij*_ of the partially-connected Ising model. Plots of the fields *h*_*i*_ (a-c), sorted by the average local activity ⟨*σ*_*i*_ ⟩^(IF)^ of the associated neuron *i*, in order of decreasing ⟨*σ*_*i*_ ⟩ ^(IF)^, and distributions of the non-zero interaction constants *J*_*ij*_ (d-f), that reproduce the respective data of the Ising model presented in Fig 13 for the three different thresholds *η* = {0.10, 0.15, 0.20}, for a fully-excitatory system at criticality of size *N* = 180.

As usual, we can also analyze how the Ising model might predict quantities that are not being constrained by the BM algorithm. In Fig 15 we present the comparison between the tree-point correlation functions *T*_*ijk*_ measured in the partially-connected Ising model and the IF model. As in the fully-connected case, the Ising model consistently overestimates the values of *T*_*ijk*_. The quality of the fit for the smallest threshold considered, *η* = 0.10, is comparable to the fully-connected Ising models (Fig 6). However, as *η* increases, the quality of the fit declines, particularly for the smallest 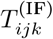. Next, we present in Fig 16 the results for the probability *P* (*K*) of simultaneous firing. The predictive capability of the partially-connected Ising model for small *K* is similar to that of fully-connected ones (Fig 7), while significant differences can be seen for large *K*. As the threshold *η* is increased, the Ising model seems to predict larger simultaneous activity at the scale of the whole network, where *K ≈ N*, indicated by a local maximum in *P* (*K*). This seems to indicate that the small *J*_*ij*_ that we are disregarding encode the information concerning the large *K* regime of *P* (*K*).

**Fig 15.**
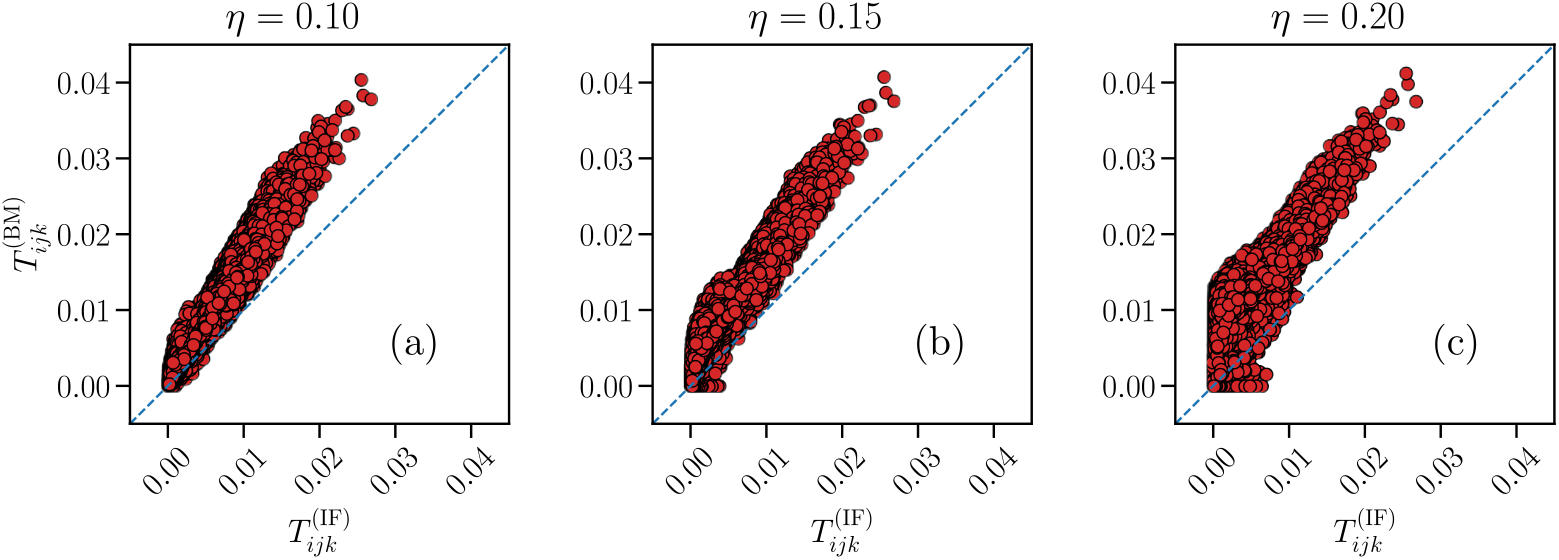
Predictive capability of the partially-connected Ising model for the three-point correlation functions *T*_*ijk*_. Comparison between the three-point correlation functions *T*_*ijk*_ of the partially-connected Ising-like model (y-axes) and the IF network (x-axes), for the three thresholds *η* = {0.10, 0.15, 0.20}, for a fully-excitatory system at criticality of size *N* = 180. Results are averages over *N*_*b*_ = 10^7^ time bins for the IF model (IF), and averages over *M*_*c*_ = 3 *·*10^7^ spin configurations for the Ising model (BM). Error bars are given by the standard error, and most are smaller or equal to the symbol size.

**Fig 16.**
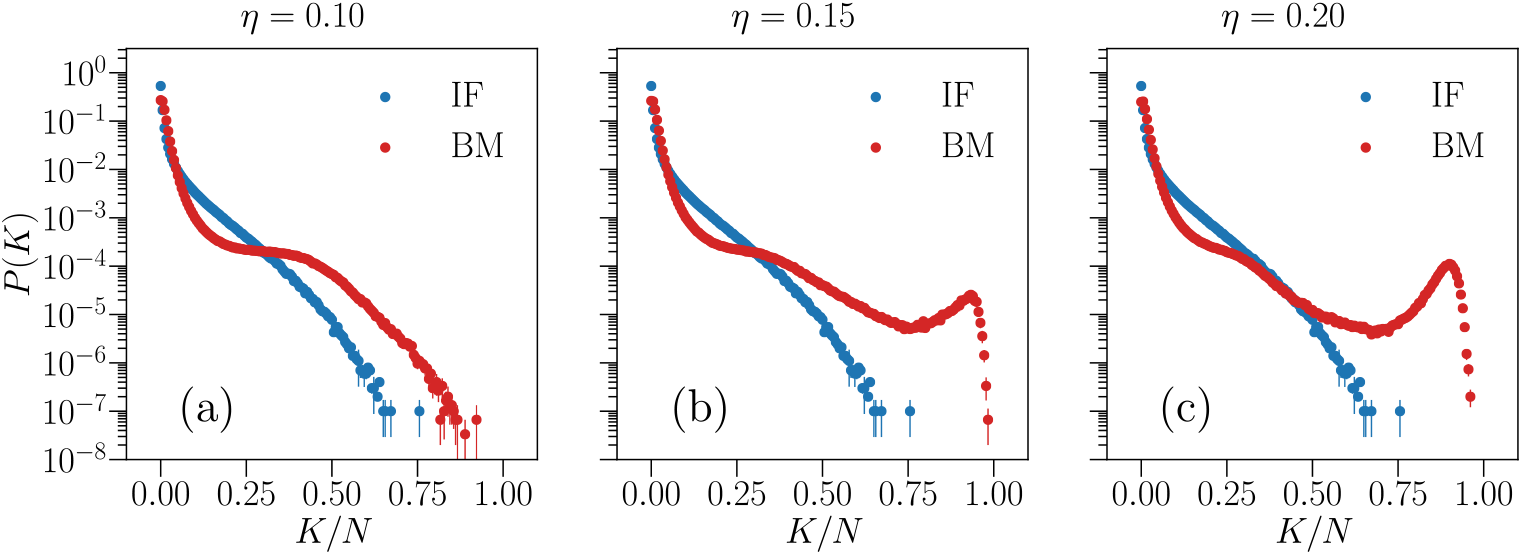
Predictive capability of the partially-connected Ising model for the probability of simultaneous firing *P* (*K*). Comparison between the simultaneous firing/up-state probability *P* (*K*) of the Ising-like model (red) and the IF model network (blue), for the three thresholds *η* = {0.10, 0.15, 0.20}, for a fully-excitatory system at criticality of size *N* = 180. Results are averages over *N*_*b*_ = 10^7^ time bins for the IF model (IF), and averages over *M*_*c*_ = 3 *·*10^7^ spin configurations for the Ising model (BM). Error bars are given by the standard error, and most are smaller or equal to the symbol size.

Finally, in Fig 17 we plot the thermodynamic functions *m, χ* and *C*_*v*_ as a function of the temperature *T* for the partially-connected Ising systems, considering the three cases *η* = {0.10, 0.15, 0.20 }. Results are quite robust with respect to *η*, taking into account that there is a difference of *≈* 31.9% of removed *J*_*ij*_ between the smallest and largest thresholds *η* considered. The tendency for the susceptibility *χ* and specific heat *C*_*v*_ to diverge with *N* is also still clearly visible when comparing to the results for a smaller system size with *N* = 120 (black symbols), which in turn considers the full set {*J*_*ij*_}.

**Fig 17.**
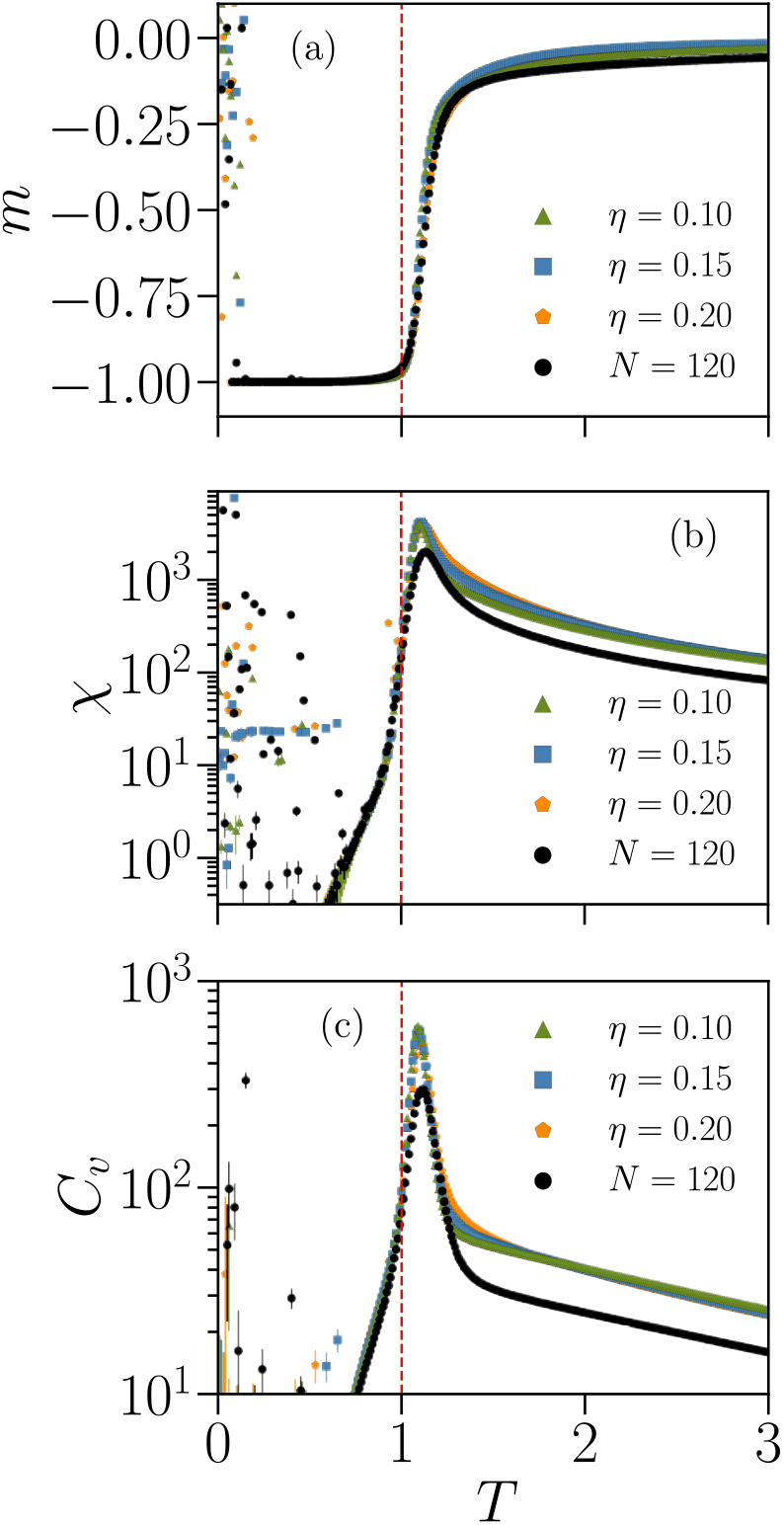
Thermodynamic functions of partially-connected Ising models associated with an IF model network with *N* = 180. Average magnetization per spin *m* (a), susceptibility *χ* (b) and specific heat *C*_*v*_ (c) as a function of the temperature *T* ∈ [0.1, 3.0], simulated using the learned parameters shown in Fig 14 for the three different thresholds *η* = ∈ {0.10, 0.15, 0.20}, for a fully-excitatory IF network at criticality of size *N* = 180. The black symbols show the results for a learned Ising model associated with an IF network of size *N* = 120, considering the full set {*J*_*ij*_ }. As in Figs 8 and 12, the cloud of random values for *T <* 1 suggests the presence of a spin-glass phase. Results are averages over *M*_*c*_ = 3 *·*10^6^ spin configurations. Error bars are given by the standard error and are overall smaller or equal to the symbol size.

## Discussion

By means of the Maximum Entropy method (MEM), we mapped the local and pairwise information of IF complex neural networks into Ising models with frustrated spins. Independently of the system size *N*, the local fields *h*_*i*_ are mostly negative (Fig 4a-d) and the distribution of interaction constants *J*_*ij*_ (Fig 4e-h) is bimodal for small systems, but tends to a normal distribution centered around zero as *N* increases. These Ising systems display a spin glass phase at low temperatures, independently of *N* (Fig 8), and the susceptibility and specific heat tend to diverge with the system size, with a maximum near *T* = *T*_0_ = 1, the effective temperature used in the BM learning to fit the IF network neuronal data. This is an indication that the Ising model analogs of the IF networks operate near a critical point. Furthermore, at least for *N ≤* 120, the height of the maximum in the susceptibility is sensible to changes in the details of the connectivity of the network (Fig 8b). We remark that the Ising models do not predict well unconstrained quantities such as the three-point correlation functions (Fig 6) and the probability of simultaneous firing even for whole networks (Fig 7). This discrepancy is enhanced when considering only a subset of the neurons (Fig 10), likely due to inputs from neurons outside the subnetwork which are not considered in the MEM mapping. This indicates that additional caution should be taken when analysing real neural networks using the MEM approach, which often consider only a subset of the total neuronal population. The presence of inhibitory neurons in the IF networks leads to reduced thermal fluctuations in the associated Ising models, evidenced by a decrease of the maxima near the critical point (Fig 12).

Ultimately, the potential of the MEM approach using a numerical model of neural activity is limited by its CPU time demand when considering large system sizes *N* ⪆ 120. To circumvent this issue, we consider a partially-connected Ising model with only a subset of the total pairs of interaction constants {*J*_*ij*_ } obtained considering only the largest correlation functions measured in the IF network, allowing to study a system of size *N* = 180 (Figs 13-17). Similar MEM procedures allow the study of much larger sizes, such as the so-called restricted Boltzmann Machine [57] or a random projection model [58], but the resulting maximum entropy distributions from these approaches are no longer analogous to that of an Ising model, and the thermodynamical interpretation is no longer applicable [27]. A very recent study [59] shows that it might be possible to consider large networks considering only connections which contain the maximum mutual information among all pairs of neurons. Since the IF model can also tune systems off the critical state, this approach could potentially be used to study thermodynamic properties of networks in the sub- and supercritical state, brain states associated with pathological conditions [60].

## Methods

### Integrate-and-fire model

The model considers *N* neurons placed randomly inside a cubic space of side 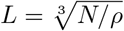, where *ρ* = 0.016 [55] is the density of the neurons. Connections are directed and weighted, with dynamic synaptic strengths *w*_*ij*_ *∈* [0, 1] between pre- and post-synaptic neurons *i* and *j*. Each neuron has at least one incoming connection, and the distribution of out-going degrees *k*_out_ *∈* [2, 20] is a power law, i.e.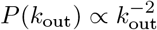, following experimental measurements in functional networks [61]. The probability that two neurons are connected decays exponentially with their euclidean distance *r, P* (*r*) *∝ e*^*−r/r*0^, where *r*_0_ = 5 is a characteristic length [62]. Neurons can be excitatory or inhibitory, with fraction *p*_in_. We implement synaptic plasticity, in which the strengths *w*_*ij*_ can change dynamically with time. We consider both short-term and long-term plasticity (STP and LTP for short, respectively). STP refers to the modification of synaptic strengths on a short timescale, of the order of milliseconds. This is mainly caused by the depletion of synaptic resources over time required for synaptic transmission [63]. We denote by *u*_*i*_(*t*)*∈* [0, 1] the normalized amount of neurotransmitters available to neuron *i* at time *t*, representing the so-called readily releasable pool of neurotransmitter vesicles [64, 65]. On the other hand, LTP is a Hebbian-like process which strengthens or weakens connections strengths *g*_*ij*_ *∈* (0, 1] depending on their usage over time, acting on much longer timescales, ranging from minutes to hours or even years [66]. We therefore separate the synaptic strengths *w*_*ij*_(*t*) = *u*_*i*_(*t*)*g*_*ij*_ into the two components *u*_*i*_(*t*) and *g*_*ij*_, to emphasize the role of these two plasticity mechanisms. The time dependence of *w*_*ij*_(*t*) comes only from the STP term *u*_*i*_(*t*), since the LTP strength terms *g*_*ij*_ can be regarded as constant compared to the timescale of the dynamics.

Each neuron *i* is characterized by a membrane potential *v*_*i*_. A neuron *i* will fire at some time *t* when its potential *v*_*i*_ surpasses a threshold *v*_*c*_ = 1. Activity will then propagate to all post-synaptic neurons *j* with incoming connections from *i* according to the following equations [31]:

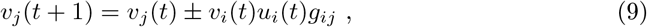

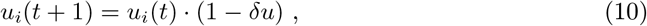

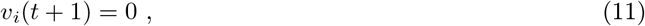

where + and *−* stands for excitatory and inhibitory pre-synaptic neuron, respectively, and *δu* = 0.05 represents the fractional amount of neurotransmitter released at each neuronal firing [67]. The timestep unit corresponds roughly to the joint interval of synaptic and axonal delay, i.e. the time interval between the generation of the action potential at the pre-synaptic neuron and the membrane potential change at the post-synaptic one, and is of the order of 10 milliseconds [68, 69]. Equation (9) models the signal transmission from pre-to post-synaptic neurons *i* and *j*, respectively. Equation (10) models the STP dynamics, while equation (11) resets the membrane potential of the pre-synaptic neuron into its resting state. We set a minimum value *v*_min_ = *−* 1 for the membrane potential of each neuron, to prevent the possibility of a *v*_*i*_ being systematically decreased to overly negative values when *p*_in_ *>* 0. After firing, a neuron enters into a refractory period of a single timestep *t*_*r*_ = 1 during which it is incapable of receiving or eliciting any activity. Equations (9)-(11) can lead to the generation of neuronal avalanches [31], periods of cascading activity separated by intervals of quiescence. Their size *S* is defined by the number of neurons which fire during the avalanche, while their duration *D* is the number of numerical timesteps it lasted. To keep the activity ongoing, we implement a small external stimulation. Namely, at every timestep, even during avalanches, a voltage input *δv* = 0.1*v*_*c*_ is added to a randomly chosen neuron. From equation (10) we see that the amount of synaptic resources gradually depletes as a neuron fires, eventually rendering it incapable of transmitting further signals. In real systems, this is counteracted by a recovery mechanism, where the readily releasable pool is slowly recharged over the span of seconds [70]. Since activity propagation is of the order of milliseconds, we assume a separation of timescales [71], and recharge simultaneously the *u*_*i*_ of all neurons by a certain amount *δu*_rec_ only at the end of every avalanche,

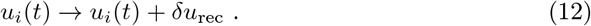

By changing the value of *δu*_rec_ one can adjust the dynamical state of the system [31, 55]. At a specific value of *δu*_rec_, the network can be tuned to a critical state, producing avalanches whose size distribution behaves as a power law with a cut-off that scales with the system size *N*. Identical systems with *δu*_rec_ below or above this tuned value would exhibit behaviour typical of sub- and supercritical systems, respectively. In a subcritical system, activity is extinguished too quickly, leading to an exponential decay in the size distribution, while for a supercritical one, an excess of large avalanches is observed, characterized instead by a local maximum of the size distribution near the cut-off [31].

Before performing any measurements, we let the dynamics evolve for a certain number of timesteps in order to shape the distribution of strengths *g*_*ij*_ by LTP. We start by setting them uniformly distributed in the interval *g*_*ij*_ *∈* [0.04, 0.06]. According to the rules of Hebbian plasticity [72–74], if some neuron *i* frequently stimulates another neuron *j*, then the synapse from *i* to *j* will be strengthened. In this case, we increase the strength of the synapses *g*_*ij*_ proportionally to the voltage variation induced in the post-synaptic neuron *j* due to *i* as,

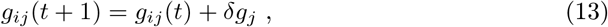

where *δg*_*j*_ = *β* |*v*_*j*_(*t* + 1) *− v*_*j*_(*t*)| and *β* = 0.04 sets the rate of this adaptation. On the other hand, synapses that are rarely active tend to weaken over time [74]. Therefore, at the end of each avalanche, we decrease all terms *g*_*ij*_ by the average increase in strength per synapse,

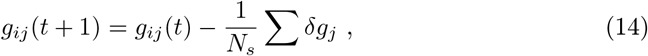

where *N*_*s*_ is the number of synapses. In real networks, synapses that weaken consistently are eventually pruned. To avoid modifying the scale-free structure of the network, we modify the strengths *g*_*ij*_ either for a fixed number *N*_aval_ = 10^4^ of avalanches or until a strength *g*_*ij*_ first reaches a minimum value *g*_min_ = 10^*−*5^, where we then set that strength to *g*_*ij*_ = *g*_min_.

### Maximum Entropy Modelling

The binarization of the IF model firing dynamics introduces the notion of the probability *P*_IF_(***σ***) to observe, during a time bin of duration Δ*t*_*b*_, any of the 2^*N*^ possible patterns of firing states ***σ*** = {*σ*_1_, *σ*_2_, …, *σ*_*N*_ }in a network of size *N*, with each *σ*_*i*_ ∈ {−1, 1 }. We are interested in defining this *P*_IF_(***σ***) in a way that is consistent with the expectation values measured in the IF model for a given network. Restating the problem more generally, we have a random variable ***σ*** defined by a probability distribution *P* (***σ***) which is unknown *a priori*. This distribution *P* (***σ***) must, of course, obey the normalization condition,

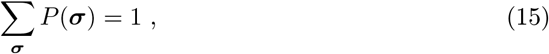

where Σ_***σ***_ indicates a sum over all possible outcomes of ***σ***. We want to find an expression for *P* (***σ***) given the average value of some function *f* (***σ***) of the random variable ***σ***,

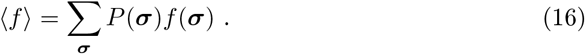

The maximum entropy principle [17] states that the best choice for *P* (***σ***) out of all possible distributions which are consistent with previous knowledge (in this case, the average ⟨*f* ⟩) is the one that has the maximum entropy, i.e. the least biased one. In other words, we should use the probability distribution that maximizes the entropy [75],

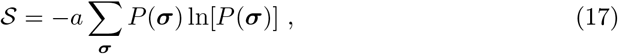

while subject to constraints (15) and (16), where *a* is a positive constant that defines the units of 𝒮, which we set to *a* = 1 without loss of generality. To solve this problem, we can make use of the method of Lagrangian multipliers [17]. For Lagrangian multipliers *λ*_0_ and *λ*_1_ associated with the constraints (15) and (16), respectively, the Lagrangian reads,

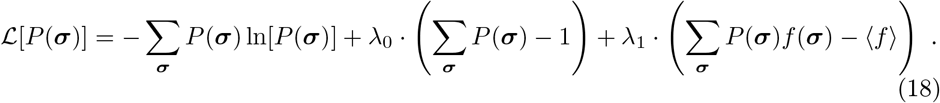

To proceed, it is useful to explicitly define each probability *p*_*i*_ *P ≡* (***σ*** = ***σ***^*i*^), where ***σ***^*i*^ is a particular outcome of the random variable ***σ***. Since the Lagrangian (18) depends on a function, *P* (***σ***), ℒ is a functional, and to search for a local extremum with respect to *P* (***σ***) we require the following functional (or variational) derivative [76],

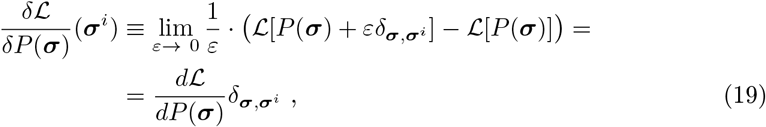

where *ε* is a real number and *δ*_***σ***,***σ****i*_ is the delta Kronecker function. The equality follows from multiplying and dividing the right side by *δ*_***σ***,***σ****i*_ and then taking the limit *ε →* 0. Equation (19) expresses the change in the functional ℒ due to changes in the function *P* (***σ***) at the particular point ***σ*** = ***σ***^*i*^. The extrema condition equation then implies

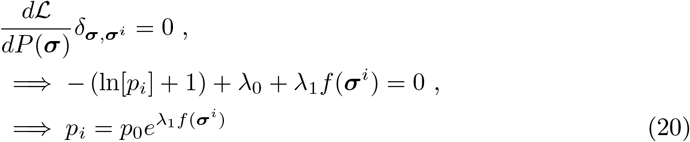

with 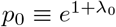. Since the last result is identical for all other probabilities *p*_*j*_ ≠_*i*_, we have for the probability distribution *P* (***σ***),

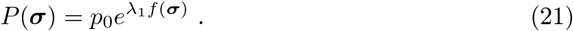

The factor *p*_0_ is determined by the normalization condition (15),

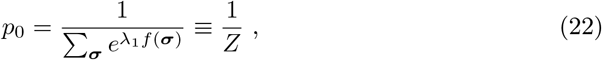

So

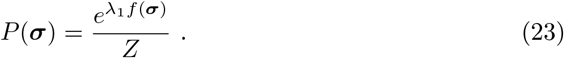

The last result can be easily generalized for the case that we have instead *C* constraints associated with the average value of *C* functions of the random variable ***σ***, { *f*_1_(***σ***), *f*_2_(***σ***), …, *f*_*C*_(***σ***) }. In this case, besides *λ*_0_, associated with the normalization condition, one has *C* Lagrangian multipliers { *λ*_1_, *λ*_2_, …, *λ*_*C*_ }, and the last term in the Lagrangian (18) needs to be modified by including a sum over the *C* constraints. This then gives

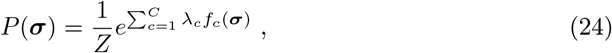

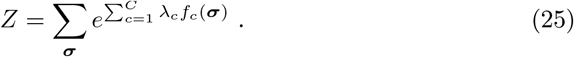

Going back now to the particular case of the IF model, for a given network with *N* neurons, as a first approximation, we can aim at constraining the single-neuron ⟨*σ*_*i*_ ⟩ and pairwise ⟨*σ*_*i*_*σ*_*j*_⟩ information, and, by extension, the correlation functions *C*_*ij*_ (see “Firing statistics” in the Results section). This corresponds to *N* + *N·* (*N−* 1)*/*2 constraints, giving *N* Lagrangian multipliers associated with each *f*_*i*_(***σ***) = *σ*_*i*_, that we will denote as *h*_*i*_, and *N ·* (*N −* 1)*/*2 multipliers for each different pair *f*_*ij*_(***σ***) = *σ*_*i*_*σ*_*j*_, which we will denote in turn as *J*_*ij*_. For this case, the maximum entropy probability (24) reads,

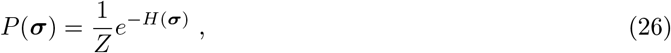

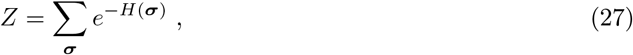

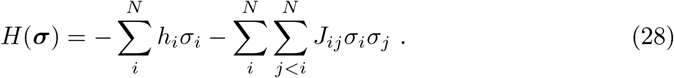

Equations (26)-(28) are the expressions for the least-biased probability distribution that is consistent with the measured values of ⟨*σ*_*i*_ ⟩ and *C*_*ij*_ in the IF model. This is the Boltzmann distribution, and is mathematically equivalent to the distribution of spin configurations of a generalized Ising model [9],

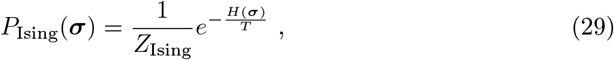

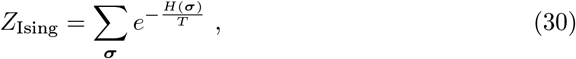

with temperature *T* = 1 in natural units (Boltzmann constant *k*_*B*_ = 1), and where *H*(***σ***) is the Hamiltonian or energy function and *Z*_Ising_ is the partition function, whose sum runs over all 2^*N*^ possible configurations of spin states ***σ***. This prompts us to interpret this maximum entropy description as a mapping from the temporal averages of the neuronal network to that of a fully-connected spin lattice. Under this conceptual view, *h*_*i*_ is analogous to a local external field acting on spin *i*, whereas *J*_*ij*_ is an interaction constant between spins *i* and *j*. The task is now to find the values of *h*_*i*_ and *J*_*ij*_ that reproduce the measured expectation values from the IF model. This is a particular example of the inverse Ising problem [77–79], also known as Boltzmann Machine (BM) learning [80], which consists in inferring the Hamiltonian of a certain complex multi-component system from its observed statistics. In principle, each parameter *h*_*i*_ and *J*_*ij*_ can be determined from the derivative of the logarithm of the partition function (27),

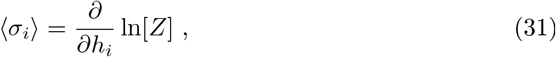

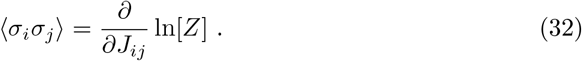

However, the number of terms in *Z* grows exponentially with *N*, as 2^*N*^, and an analytical approach becomes intractable when *N* ≳ 20. To proceed, notice that we are trying to describe an empirical distribution *P*_IF_(***σ***), as observed from the measured temporal averages of the IF model, using the analytical description *P* (***σ***), as given by equation (26). We want to choose the {*h*_*i*_} and {*J*_*ij*_} that minimize the loss in information when using *P* (***σ***) as a proxy for *P*_IF_(***σ***). Specifically, this means we want the set of {*h*_*i*_ } and {*J*_*ij*_ } that minimize the so-called Kullback-Leibler divergence [81] between these distributions,

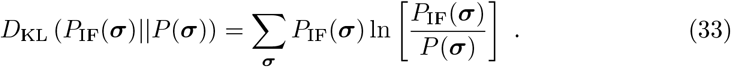

From the minimum condition equations one can show that,

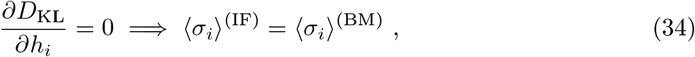

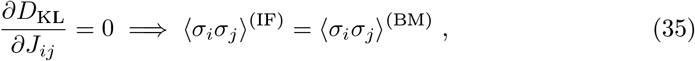

where ⟨*σ*_*i*_⟩^(IF)^ *≡* Σ_***σ***_ *σ*_*i*_*P*_IF_(***σ***) are the empirical averages measured in the IF model and and ⟨*σ* _*i*_⟩ ^(BM)^*≡* Σ _***σ***_ *σ*_*i*_ *P* (***σ***) are the ones estimated from the Boltzmann distribution (26), and analogously for the ⟨*σ*_*i*_*σ*_*j*_⟩. Therefore, a suitable method to search for the fields *h*_*i*_ and interaction constants *J*_*ij*_ of the Hamiltonian (28) is the following iterative scheme [9],

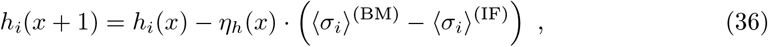

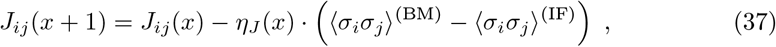

where *η*_*h*_(*x*) = 2*η*_*J*_ (*x*) *∝x*^*−α*^ are decreasing learning rates, with *α* = 0.4 for *N ≤* 40, *α* = 0.6 if 40 *< N ≤* 120, and *α* = 1.0 otherwise. We set a smaller learning rate for the {*J*_*ij*_ } since their number (*∼ N* ^2^) is much larger when compared to the {*h*_*i*_} (*N*), so we update their values at a slower rate to avoid divergences during the learning procedure. At each iteration *x*, the data sets {⟨*σ*_*i*_⟩ ^(IF)^ } and {⟨ *σ*_*i*_*σ*_*j*_ ⟩ }^(IF)^ generated by the IF model are compared to those estimated by sampling the distribution (26), {⟨*σ*_*i*_ ⟩ } ^(BM)^ and {⟨*σ*_*i*_*σ*_*j*_ ⟩ ^(BM)^ }, using the sets of fields {*h*_*i*_(*x*)} and coupling constants {*J*_*ij*_(*x*)}. We start with *h*_*i*_(*x* = 1) = ⟨*σ*_*i*_⟩ ^(IF)^ and *J*_*ij*_(*x* = 1) = 0 and then iterate equations (36) and (37) typically until *x* = *N*_BM_ ∼ 60000. At each iteration, {⟨*σ*_*i*_⟩ ^(BM)^ } and {⟨*σ*_*i*_*σ*_*j*_⟩ ^(BM)^ } are estimated from Monte Carlo simulations, using the Metropolis algorithm [82, 83], by averaging over *M*_*c*_ = 3 *·* 10^5^ spin configurations. We disregard the first 150*N* configurations for systems with *N ≤* 120, or up to 10^5^*N* ^2^ configurations for the *N* = 180 case, in order to reduce correlations with the initial state, and use only every 2*N* -th configuration for averaging, to reduce autocorrelations. At the end of the learning routine, we study the generalized Ising models with the set of fitted parameters {*h*_*i*_ }and {*J*_*ij*_ } by sampling the distribution (26), averaging over an increased amount of spin configurations *M*_*c*_ = 3 *·* 10^6^ for systems with *N ≤* 120, or up to *M*_*c*_ = 3 *·* 10^7^ configurations for the system with *N* = 180, to reduce error bars.

## Acknowledgements

L.d.A. acknowledges support from the Italian MUR project PRIN2017WZFTZP and from NEXTGENERATIONEU (NGEU) funded by the Ministry of University and Research (MUR), National Recovery and Resilience Plan (NRRP), and project MNESYS (PE0000006)— A multiscale integrated approach to the study of the nervous system in health and disease (DN. 1553 11.10.2022).

## Supporting information

**S1 Fig. Avalanche size and duration distributions of critical fully-excitatory IF networks**.

**S1 Table. Values of** *δu*_**rec**_ **for the fully-excitatory systems considered in S1 Fig**

**S2 Fig. Avalanche size and duration distributions of critical IF networks with different fractions of inhibitory neurons**.

**S2 Table. Values of** *δu*_**rec**_ **for the systems with** *N* = 80 **considered in S2 Fig**

**S3 Fig. Dependence with the initial spin configuration of the thermodynamic functions of Ising-like models associated with fully-excitatory IF networks**.

**S4 Fig. Quality test for the BM learning process for IF subnetworks**.

**S5 Fig. Quality test for the BM learning process for IF subnetworks with different fractions of inhibitory neurons**.

**S6 Fig. Learned fields** *h*_*i*_ **of the pairwise Ising model associated with an IF network with inhibitory neurons**.

**S7 Fig. Quality test for the BM learning process with a partially-connected Ising model of the average local activities**.

## Notes

### Competing Interest Statement

The authors have declared no competing interest.

### Summary of Updates

Added supplementary information

