## Supplementary information for "Thermodynamic analog of integrate-and-fire neuronal networks by maximum entropy modelling"

(Dated: January 13, 2024)

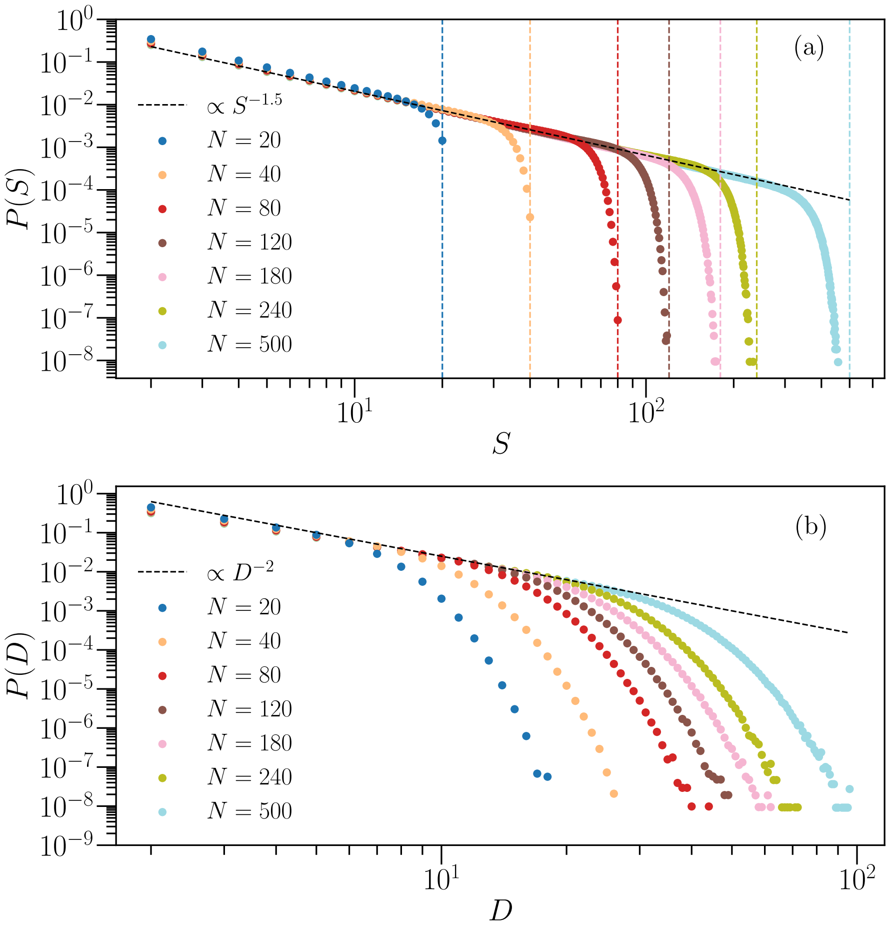


FIG. S1. **Avalanche size and duration distributions of critical fully-excitatory IF networks.** Avalanche size $S$ (a) and duration $D$ (b) distributions for several system sizes $N$ tuned to the critical state. The values of $\delta u_{\rm rec}$ for each $N$ are reported in table (S1). Results are averages over $2 \cdot 10^8$ avalanches, generated over $N_c = 10^4$ different network configurations. The vertical dashed lines in (a) indicate the system size of the corresponding colour-matching curve of $P(S)$.

TABLE S1. Values of $\delta u_{\rm rec}$ for the fully-excitatory systems considered in Fig. (S1)

| **System size** $N$ | $\delta u_{\bf rec}$ |
|---|---|
| 20 | 0.00190 |
| 40 | 0.00150 |
| 80 | 0.00120 |
| 120 | 0.00100 |
| 180 | 0.00080 |
| 240 | 0.00075 |
| 500 | 0.00050 |

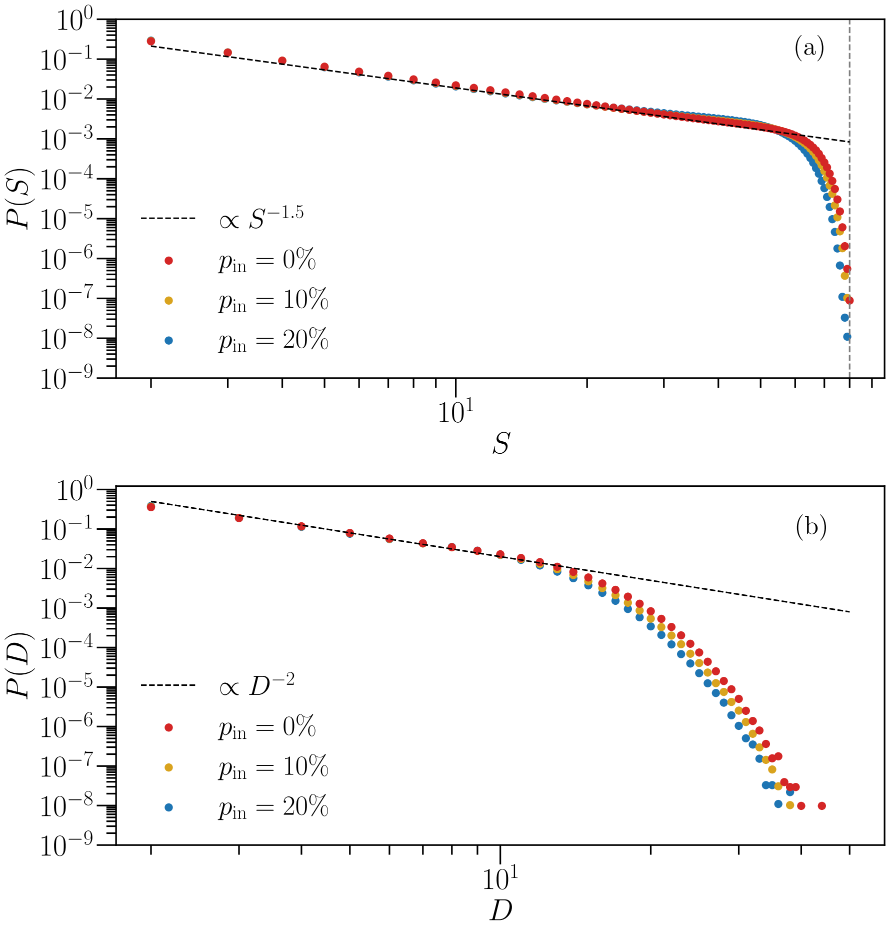


FIG. S2. **Avalanche size and duration distributions of critical IF networks with different fractions of inhibitory neurons.** Avalanche size $S$ (a) and duration $D$ (b) distributions for systems with $N = 80$ and different percentage $p_{\rm in}$ of inhibitory neurons, tuned to the critical state. The values of $\delta u_{\rm rec}$ for each $p_{\rm in}$ are reported in table (S2). Results are averages over $2 \cdot 10^8$ avalanches, generated over $N_c = 10^4$ different network configurations. The vertical dashed line in (a) indicates the system size $N = 80$.

TABLE S2. Values of $\delta u_{\rm rec}$ for the systems with $N = 80$ considered in Fig. (S2)

| $p_{\bf in}$ | $\delta u_{\bf rec}$ |
|---|---|
| 0% | 0.0012 |
| 10% | 0.0014 |
| 20% | 0.0017 |

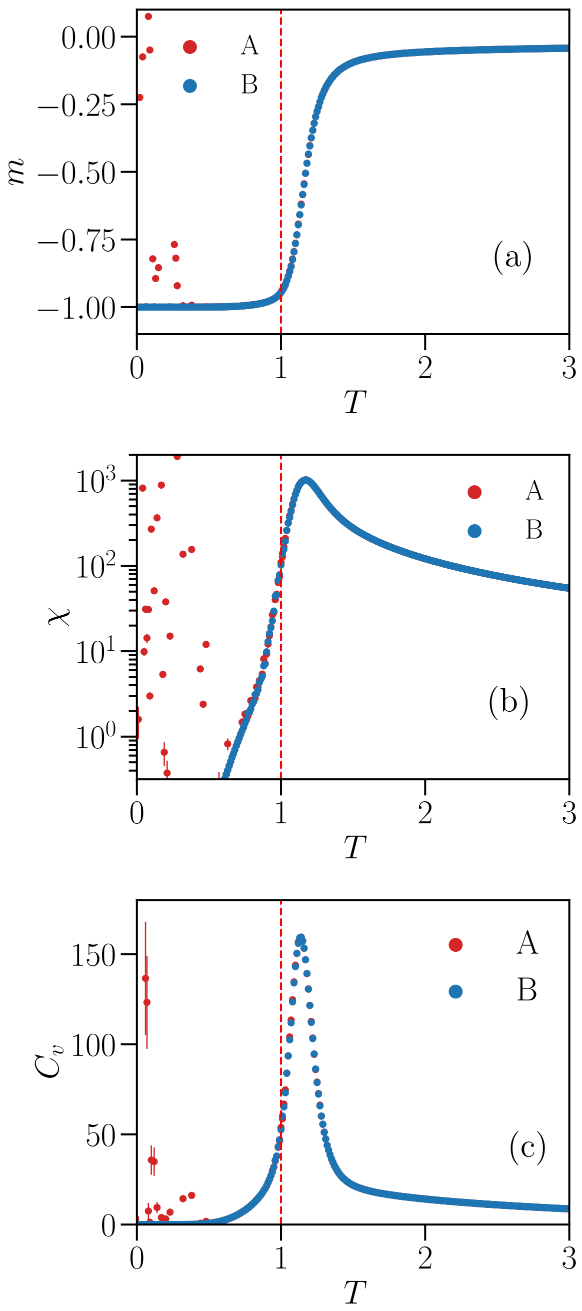


FIG. S3. **Dependence with the initial spin configuration of the thermodynamic functions of Ising-like models associated with fully-excitatory IF networks.** Magnetization per spin $m$ (a), susceptibility $\chi$ (b) and specific heat $C_v$ (c) as a function of the temperature $T$ of a pairwise Ising system associated to a network with $N = 80$ neurons, using two different initial spin configurations for MC sampling: starting with random $\sigma_i$ (A) or with all $\sigma_i = -1$ (B). Results are averages over $M_c = 3 \cdot 10^6$ spin configurations. Error bars are given by the standard error.

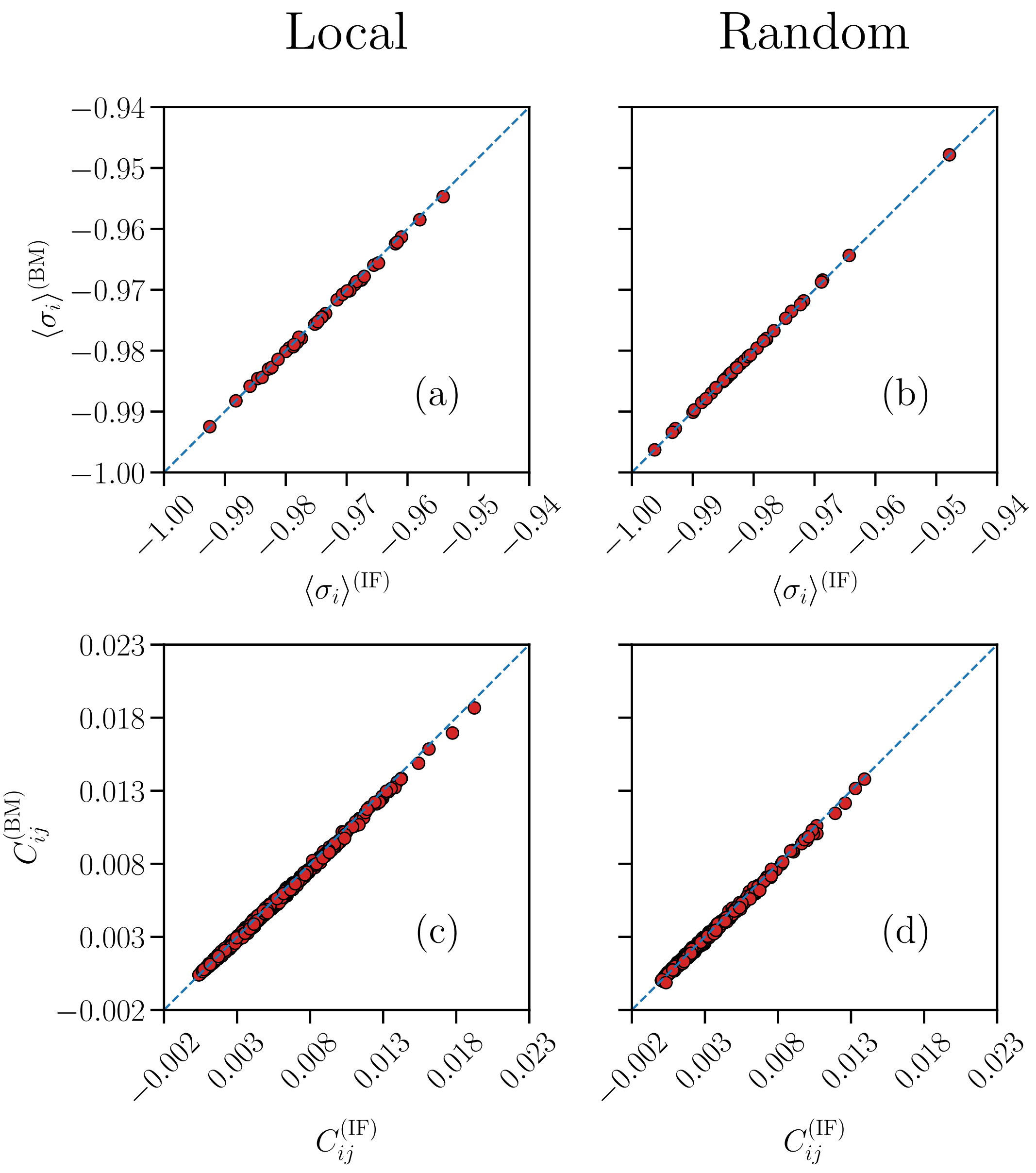


FIG. S4. **Quality test for the BM learning process for IF subnetworks.** Comparison between the average local activity $\langle\sigma_i\rangle$ (a-b) and correlation functions $C_{ij}$ (c-d) of the pairwise Ising model ($y$-axes) and IF model network ($x$-axes), for subnetworks with $n = 40$ neurons, with a local and random spatial distribution, in a system with $N = 500$ neurons. The blue dashed lines are given by the bisector $y = x$. Results are averages over $N_b = 10^7$ time bins for the IF model (IF), and averages over $M_c = 3 \cdot 10^6$ spin configurations for the Ising model (BM). Error bars are given by the standard error, and are smaller or equal to the symbol size.

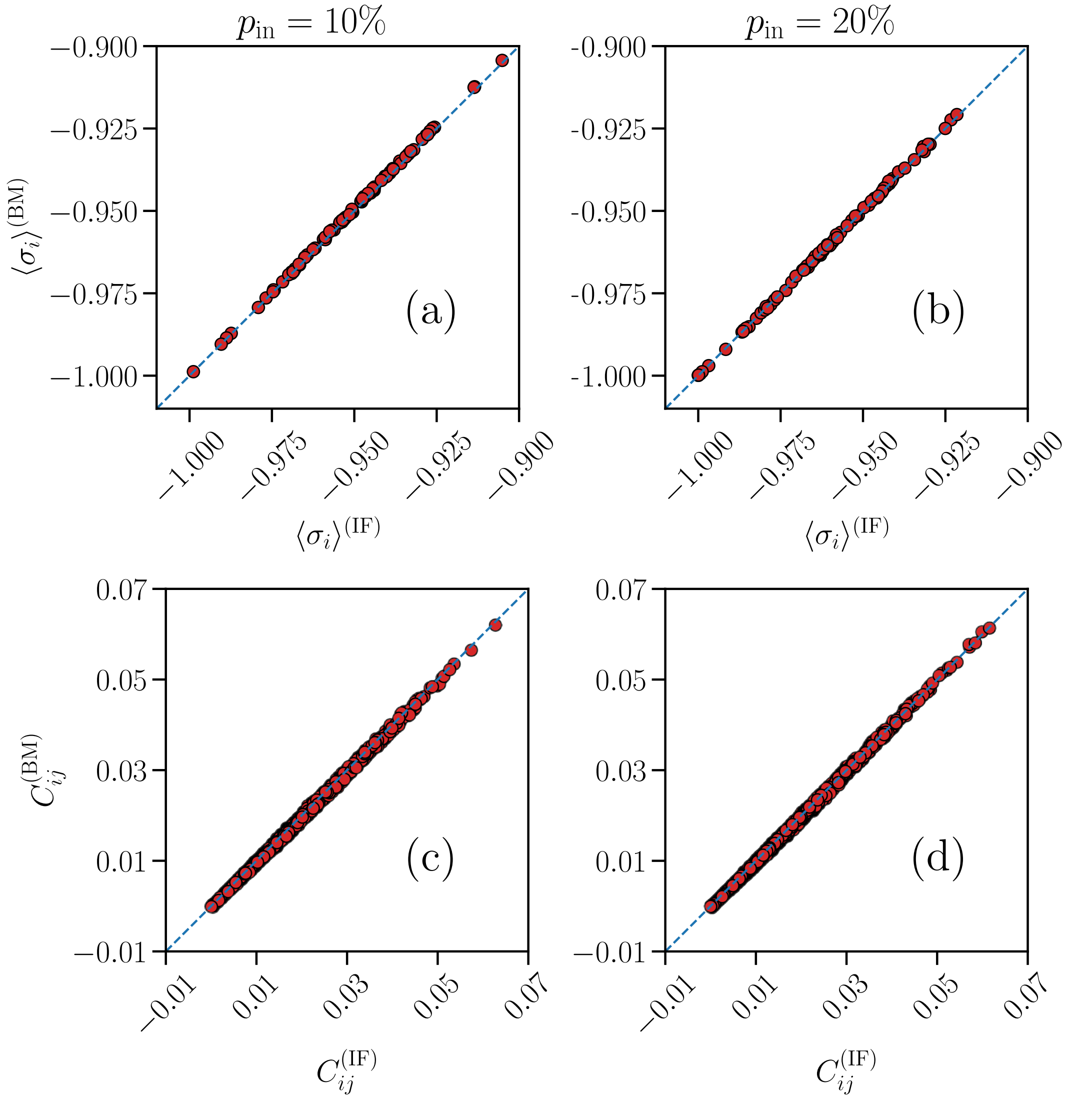


FIG. S5. **Quality test for the BM learning process for IF subnetworks with different fractions of inhibitory neurons.** Comparison between the average local activity $\langle\sigma_i\rangle$ (a-b) and correlation functions $C_{ij}$ (c-d) of the pairwise Ising model ($y$-axes) and IF model network ($x$-axes), for a IF network of size $N = 80$ with different fractions $p_{\rm in} = \{10\%, 20\%\}$ of inhibitory neurons. The blue dashed lines are given by the bisector $y = x$. Results are averages over $N_b = 10^7$ time bins for the IF model (IF), and averages over $M_c = 3 \cdot 10^6$ spin configurations for the Ising model (BM). Error bars are given by the standard error, and are smaller or equal to the symbol size.

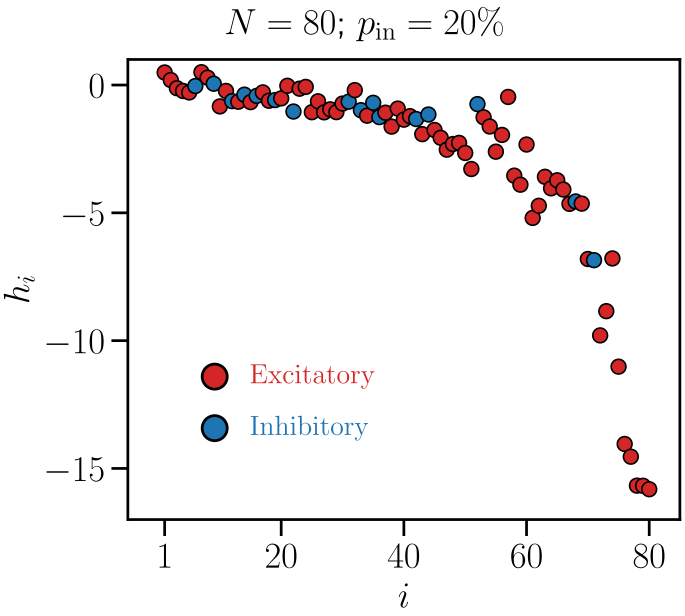


FIG. S6. **Learned fields $h_i$ of the pairwise Ising model associated with an IF network with inhibitory neurons.** Plot of the fields $h_i$, sorted by the average local activity $\langle \sigma_i \rangle^{(\rm IF)}$ of the associated neuron $i$, in order of decreasing $\langle \sigma_i \rangle^{(\rm IF)}$, for a IF network with $N = 80$ and $p_{\rm in} = 20\%$ inhibitory neurons (same data of the distribution presented in Fig. (11)a in the main text, for the same $p_{\rm in}$). Red symbols indicate fields $h_i$ associated with an excitatory neuron, while blue symbols correspond to $h_i$ associated with inhibitory ones.

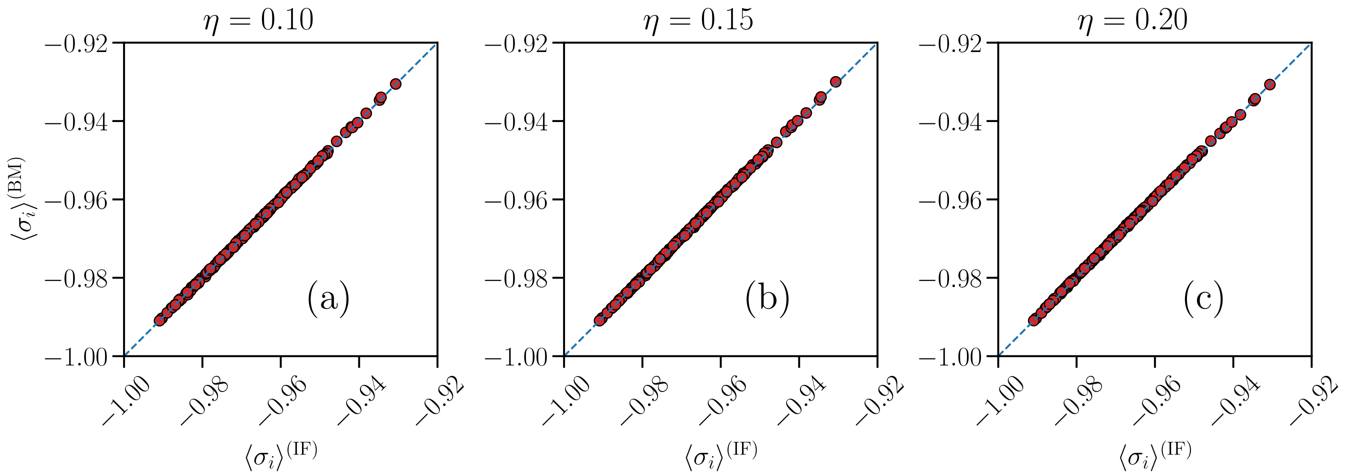


FIG. S7. **Quality test for the BM learning process with a partially-connected Ising model of the average local activities.** Comparison between the average local activity $\langle \sigma_i \rangle$ of the partially-connected pairwise Ising model ($y$-axes) and IF model network ($x$-axes), for a fully-excitatory system of size $N = 180$, considering three different thresholds $\eta = \{0.10, 0.15, 0.20\}$ for the removal of a subset of the $J_{ij}$ (see main text). The blue dashed lines are given by the bisector $y = x$. Results are averages over $N_b = 10^7$ time bins for the IF model (IF), and averages over $M_c = 3 \cdot 10^7$ spin configurations for the Ising model (BM). Error bars are given by the standard error, and are smaller or equal to the symbol size.
